# Multiple regulatory mechanisms control the production of CmrRST, an atypical signal transduction system in *Clostridioides difficile*

**DOI:** 10.1101/2021.10.06.463453

**Authors:** Elizabeth M. Garrett, Anchal Mehra, Ognjen Sekulovic, Rita Tamayo

**Affiliations:** Department of Microbiology and Immunology, University of North Carolina Chapel Hill, 125 Mason Farm Road CB#7290, Chapel Hill, NC, 27599, USA; Department of Molecular Biology and Microbiology, Tufts University School of Medicine, 136 Harrison Avenue, Boston, MA, 02111, USA

**Keywords:** Phase variation, phenotypic heterogeneity, bet-hedging, cyclic diguanylate, c-di-GMP, motility

## Abstract

*Clostridioides difficile*, an intestinal pathogen and leading cause of nosocomial infection, exhibits extensive phenotypic heterogeneity through phase variation by site-specific recombination. The signal transduction system CmrRST, which encodes two response regulators (CmrR and CmrT) and a sensor kinase (CmrS), impacts *C. difficile* cell and colony morphology, surface and swimming motility, biofilm formation, and virulence in an animal model. CmrRST is subject to phase variation through site-specific recombination and reversible inversion of the ‘*cmr* switch’, and expression of *cmrRST* is also regulated by c-di-GMP through a riboswitch. The goal of this study was to determine how the *cmr* switch and c-di-GMP work together to regulate *cmrRST* expression. We generated “phase locked” strains by mutating key residues in the right inverted repeat flanking the *cmr* switch. Phenotypic characterization of these phase locked *cmr*-ON and -OFF strains demonstrates that they cannot switch between rough and smooth colony morphologies, respectively, or other CmrRST-associated phenotypes. Manipulation of c-di-GMP levels in these mutants showed that c-di-GMP promotes *cmrRST* expression and associated phenotypes independent of *cmr* switch orientation. We identified multiple promoters controlling *cmrRST* transcription, including one within the ON orientation of *cmr* switch and another that is positively autoregulated by CmrR. Overall, this work reveals a complex regulatory network that governs *cmrRST* expression and a unique intersection of phase variation and c-di-GMP signaling. These findings suggest that multiple environmental signals impact the production of this signaling transduction system.

**IMPORTANCE:** *Clostridioides difficile* is a leading cause of hospital-acquired intestinal infections in the U.S. The CmrRST signal transduction system controls numerous physiological traits and processes in *C. difficile*, including cell and colony morphology, motility, biofilm formation, and virulence. Here we define the complex, multi-level regulation of *cmrRST* expression, including stochastic control through phase variation, modulation by the second messenger c-di-GMP, and positive autoregulation by CmrR. The results of this study suggest that multiple, distinct environmental stimuli and selective pressures must be integrated to appropriately control *cmrRST* expression.

## INTRODUCTION

The ability to adapt to environmental changes is critical to bacterial survival, including that of pathogens, which can face rapidly changing conditions and stresses during infection (1–4). Sense-and-respond adaptation strategies often involve two-component systems (TCS) that consist of a sensor kinase and a cognate response regulator. In response to an activating signal, such as binding of a ligand or a change in pH, the sensor kinase autophosphorylates and activates the response regulator through transfer of the phosphoryl group (5,6). Response regulators have a wide range of functions, and many control transcription through direct binding of DNA. The resulting transcriptional changes contribute to adaptation of the bacterium to the environmental stimulus sensed by the sensor kinase. Bacteria can also adapt to stimuli via intracellular small molecules such as cyclic diguanylate (c-di-GMP) (7,8). The intracellular level of c-di-GMP is modulated by the opposing activities of diguanylate cyclases and phosphodiesterases that synthesize and degrade c-di-GMP, respectively; the production and function of these enzymes is controlled by environmental signals (9,10). C-di-GMP is then recognized by specific protein or RNA receptors (riboswitches) that mediate the adaptive response (11–13).

In contrast, diversification of phenotypes in a bacterial population serves as a bet-hedging strategy to help ensure survival of the population as a whole. The development of phenotypically distinct variants, independent of environmental conditions, improves the odds that a subpopulation survives a sudden stress (2,14). Phase variation, a mechanism of generating phenotypic heterogeneity, occurs through reversible genetic changes that typically cause an ON/OFF phenotypic “switch” (15,16). Several mechanisms of phase variation have been described including conservative site-specific recombination, in which a sequence-specific recombinase binds inverted repeats and mediates inversion of the intervening DNA. The invertible DNA element contains regulatory information, such as a promoter, that impacts the expression of adjacent genes. In a well-characterized example in *Escherichia coli*, phase variation of fimbriae production is mediated by the *fimS* invertible element, which contains a promoter that drives transcription of the fimbrial genes when properly oriented (17–19). Additional mechanisms have been recently described. Phase variation of the cell wall protein CwpV in *Clostridioides difficile* occurs as a result of site-specific recombination of a sequence mapping to the 5’ untranslated region of *cwpV*. In one orientation, the invertible DNA sequence results in the formation of an intrinsic terminator in the mRNA when in one orientation, preventing expression of the downstream gene *cwpV*; the intrinsic terminator does not form in the mRNA with the sequence in the inverse orientation, allowing *cwpV* transcription to occur (20). Phase variation of flagellum and toxin biosynthesis in *C. difficile* also occurs through a mRNA-mediated mechanism, where one orientation of the invertible sequence yields an mRNA permissive for transcriptional readthrough, while the other orientation results in Rho-mediated transcription termination (21–24).

*Clostridioides difficile* is an intestinal pathogen and a leading cause of nosocomial infections in the U.S. *C. difficile* infection (CDI) can result in mild to severe diarrhea and potentially fatal complications such as pseudomembranous colitis, toxic megacolon, and sepsis. Recent work has shown that *C. difficile* contains multiple invertible DNA elements flanked by inverted repeats (including those noted above), indicating a considerable capacity for phenotypic heterogeneity through phase variation (25,26). Three of these invertible elements have been demonstrated to regulate downstream genes and related phenotypes in a phase variable manner. The Cdi4 invertible element, also called the flagellar switch, modulates expression of the *flgB* flagellar operon resulting in phase variation of flagella (21,22). This operon encodes the sigma factor SigD, which promotes the transcription of flagellar genes as well as transcription of *tcdR*, which encodes a direct activator of the *C. difficile* toxin genes *tcdA* and *tcdB* (27,28). Accordingly, the production of these toxins is also phase variable (21–23). The Cdi1 invertible element mediates phase variation of CwpV, which contributes to phage resistance (20,29–31). Finally, the Cdi6 invertible element, here called the “*cmr* switch”, regulates the expression of *cmrRST* in a phase variable manner (25,32). The *cmrRST* operon encodes a putative non-canonical TCS with two DNA-binding response regulators (CmrR and CmrT) and a sensor kinase (CmrS) (32). Both CmrR and CmrT contain phosphoreceiver and DNA binding domains suggesting that they function as transcriptional regulators. Phase variation of CmrRST allows *C. difficile* to switch between rough and smooth colony morphologies that differ in several physiological characteristics. Through unknown mechanisms, CmrRST positively regulates type IV pilus-independent surface motility and cell elongation, and it negatively regulates swimming motility and biofilm formation. Furthermore, a *cmrR* mutant strain is deficient for colonization and shows attenuated virulence in a hamster model of infection, indicating a role for this regulatory system in CDI.

C-di-GMP riboswitches are widespread in the *C. difficile* genome and appear to be the primary mechanism of c-di-GMP regulation in *C. difficile* (11,12,33,34). Upstream of the *cmr* switch are encoded a σ^A^-dependent promoter followed by a class II c-di-GMP riboswitch that positively regulates *cmrRST* transcription in response to c-di-GMP (Figure 1A) (33,34). Increasing c-di-GMP levels results in the formation of the rough colony morphology, consistent with increased *cmrRST* expression (32). The relative contributions of the c-di-GMP riboswitch and the *cmr* switch to controlling expression of *cmrRST*, and therefore to the phenotypes controlled by this system, are unknown.

**Figure 1.**
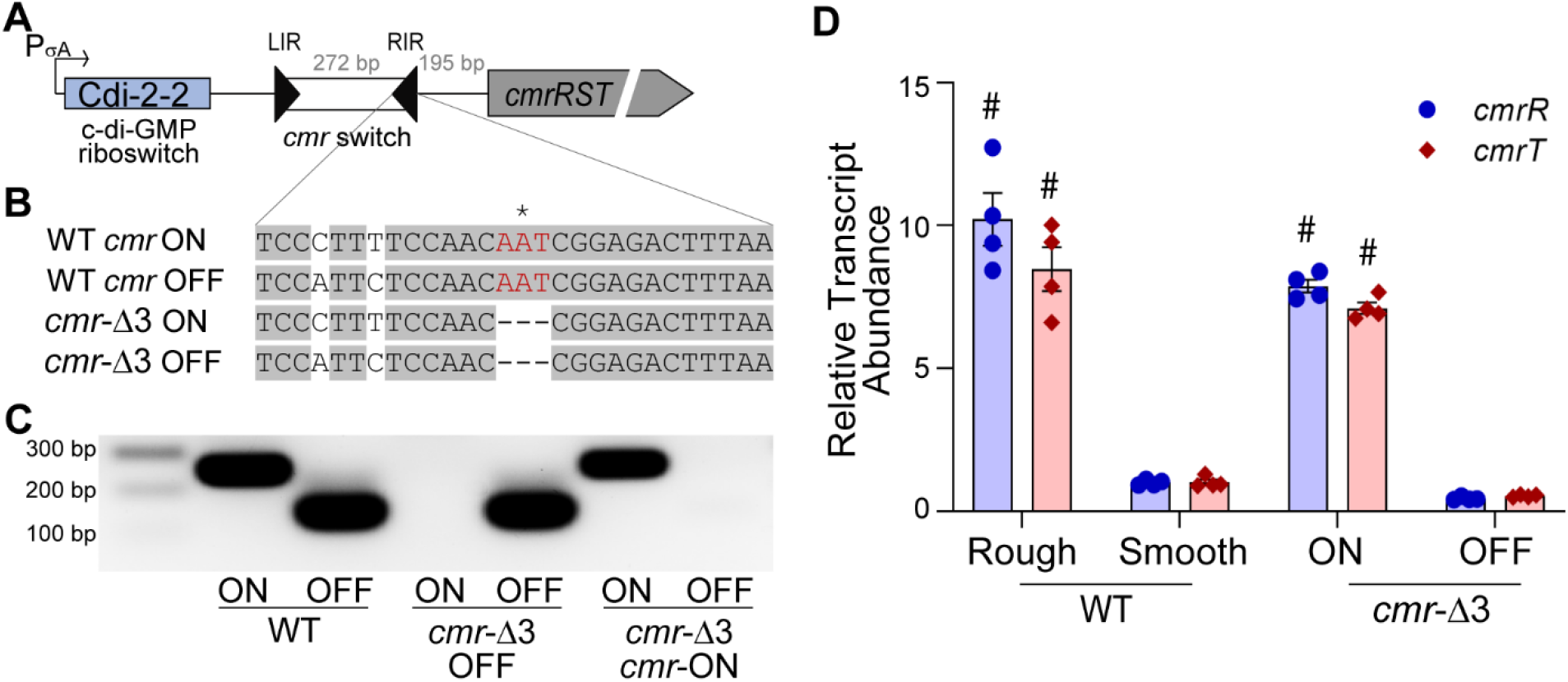
Generation of genetically phase-locked strains. (A) Diagram of the *cmrRST* locus. The previously identified σ^A^-dependent promoter and c-di-GMP riboswitch Cdi-2-2 are indicated. Black triangles represent the inverted repeats flanking the invertible DNA sequence (*cmr* switch). The OFF orientation corresponds to the sequence present in the R20291 reference genome (FN545816); the ON orientation corresponds to the inverse of that sequence. (B) Sequence of the right inverted repeat (RIR) of the *cmr* switch. Gray shading indicates sequence identity between the imperfect inverted repeats of the RIR in each orientation. Three nucleotides (shown in red) were deleted from the RIR to lock the *cmr* switch in the ON and OFF orientations. The asterisk (*) indicates the nucleotide at the site of recombination. (C) Orientation-specific PCR to detect each orientation of the *cmr* switch in WT, *cmr*-Δ3 OFF, and *cmr*-Δ3 ON strains. (D) qRT-PCR analysis for the *cmrR* and *cmrT* transcripts in WT rough and smooth isolates, *cmr*-Δ3 OFF and *cmr*-Δ3 ON. Data from four biological replicates were analyzed using the ΔΔCt method with *rpoC* as the reference gene and normalization to the WT smooth samples. Shown are means and standard error. #p < 0.0001, two-way ANOVA with Sidak’s multiple comparisons post-test comparing to the WT/smooth condition.

One challenge to the study of phase variation is its stochastic nature, which adds uncontrolled variation into an otherwise controlled experiment. Phase locked strains in which the invertible element is prevented from inverting can be a useful tool with which to study phase variable systems (21,35). Our previous work on CmrRST relied primarily on characterization of wild-type (WT) *C. difficile* rough and smooth colony isolates, which have a strong bias for the ON and OFF *cmr* switch orientations, respectively, but remain capable of switch inversion and phenotypic switching (32). In this study, we characterized *cmrRST* regulation through the interplay of c-di-GMP and the *cmr* switch by generating phase-locked *cmr* OFF and *cmr* ON mutants. Through phenotypic characterization of these phase locked strains and analysis of the effects of c-di-GMP and *cmr* switch orientation, we found that that these regulatory features act independently. We determined that phase variable regulation occurs through inversion of a promoter encoded in the *cmr* switch. In addition, we demonstrate that CmrR positively autoregulates expression of *cmrRST* via an additional promoter located between the *cmr* switch and the *cmrR* start codon. The results of this study indicate that *cmrRST* expression is subject to regulation both through sense-and-respond mechanisms and phase variation, highlighting the potential importance of this system to *C. difficile* physiology through the variety of activating signals.

## RESULTS

### Generation and phenotypic characterization of phase locked *cmr* ON and *cmr* OFF strains

To characterize the dual regulation of the *cmrRST* system by phase variation and c-di-GMP, we first generated phase- locked strains with the *cmr* switch fixed in the ON or OFF orientation. Phase locking can be achieved in multiple ways. One strategy is to delete the site-specific recombinase responsible for inversion. In *C. difficile* R20291, the RecV recombinase is required for inversion of the *cmr* switch (25). However, RecV mediates inversion of multiple sequences, thus a mutation in *recV* has pleiotropic effects (20,21,25). Instead, we chose to phase lock the *cmr* switch by mutating an inverted repeat sequence to prevent site-specific recombination from occurring (36). The exact nucleotides at which recombination and inversion of the *cmr* switch occurs were previously identified (25). We used allelic exchange to delete the nucleotide at the site of inversion in the right inverted repeat (RIR) (position 3,736,177) in the R20291 genome, as well as one additional nucleotide on each side (Figure 1B). Prior work determined that mutating the equivalent residues is the flagellar switch RIR eliminated switch inversion resulting on phase-locked strains (37). We recovered independent mutants with the *cmr* switch in either orientation and designated them *cmr*-Δ3 ON and *cmr-*Δ3 OFF. These mutants were subjected to orientation specific PCR to confirm that they are genotypically phase locked. In contrast to the WT parent that yielded both *cmr* ON and OFF products indicating heterogeneity in *cmr* switch orientation in that population of cells, *cmr*-Δ3 ON exclusively yielded a PCR product corresponding to the ON orientation, and *cmr*-Δ3 OFF only yielded the OFF orientation product (Figure 1C).

Previous work showed that R20291 rough and smooth colony variants correlate with *cmr* ON and OFF gene expression and phenotypes, respectively (32). Specifically, the *cmr* ON, rough colony variant exhibits greater surface growth and reduced swimming motility compared to the *cmr* OFF, smooth colony variant. Furthermore, ectopic expression of *cmrR* and *cmrT* individually stimulated rough colony formation and surface motility while inhibiting flagellum-mediated swimming motility, although only *cmrT* was required for rough colony development and surface motility. To conclusively determine the effect of *cmr* switch orientation on gene expression, we measured the abundance of the *cmrR* and *cmrT* transcripts in the *cmr*-Δ3 ON and *cmr*-Δ3 OFF mutants. Non-locked rough and smooth isolates were included as controls (32). Both *cmrR* and *cmrT* transcripts were 7- to 10-fold more abundant in *C. difficile* with the *cmr* switch in the ON orientation, whether in the naturally arising rough colony variant or in *cmr*-Δ3 ON (Figure 1D). Therefore, the three nucleotides at the site of RecV-mediated recombination are required for *cmr* switch inversion, and the ON orientation corresponding to the inverse of the published R20291 genome promotes expression of *cmrRST.*

The *cmr*-Δ3 phase-locked mutants were then evaluated for the phenotypes previously associated with *cmrRST* expression. As a control we included WT R20291, which is capable of CmrRST phase variation and generation of both rough and smooth colonies, as well as a Δ*cmrR*Δ*cmrT* double mutant that forms only smooth colonies (Figure 2A). The *cmr*-Δ3 OFF mutant formed exclusively smooth colonies, similar to Δ*cmrR*Δ*cmrT. cmr*-Δ3 ON formed rough colonies that resemble those in WT, though they appeared to have less defined topology (Figure 2A). Also consistent with increased *cmrRST* expression, *cmr*-Δ3 ON displayed greater surface motility than WT, while *cmr*-Δ3 OFF had decreased surface motility more similar to that of Δ*cmrR*Δ*cmrT* (Figure 2B, C). Finally, *cmr*-Δ3 ON showed decreased swimming motility and biofilm formation compared to all other strains (Figure 2D, E). Together, these results reinforce the model that the *cmr* switch orientation modulates expression of *cmrRST* and the assignment of the respective ON and OFF switch orientations.

**Figure 2.**
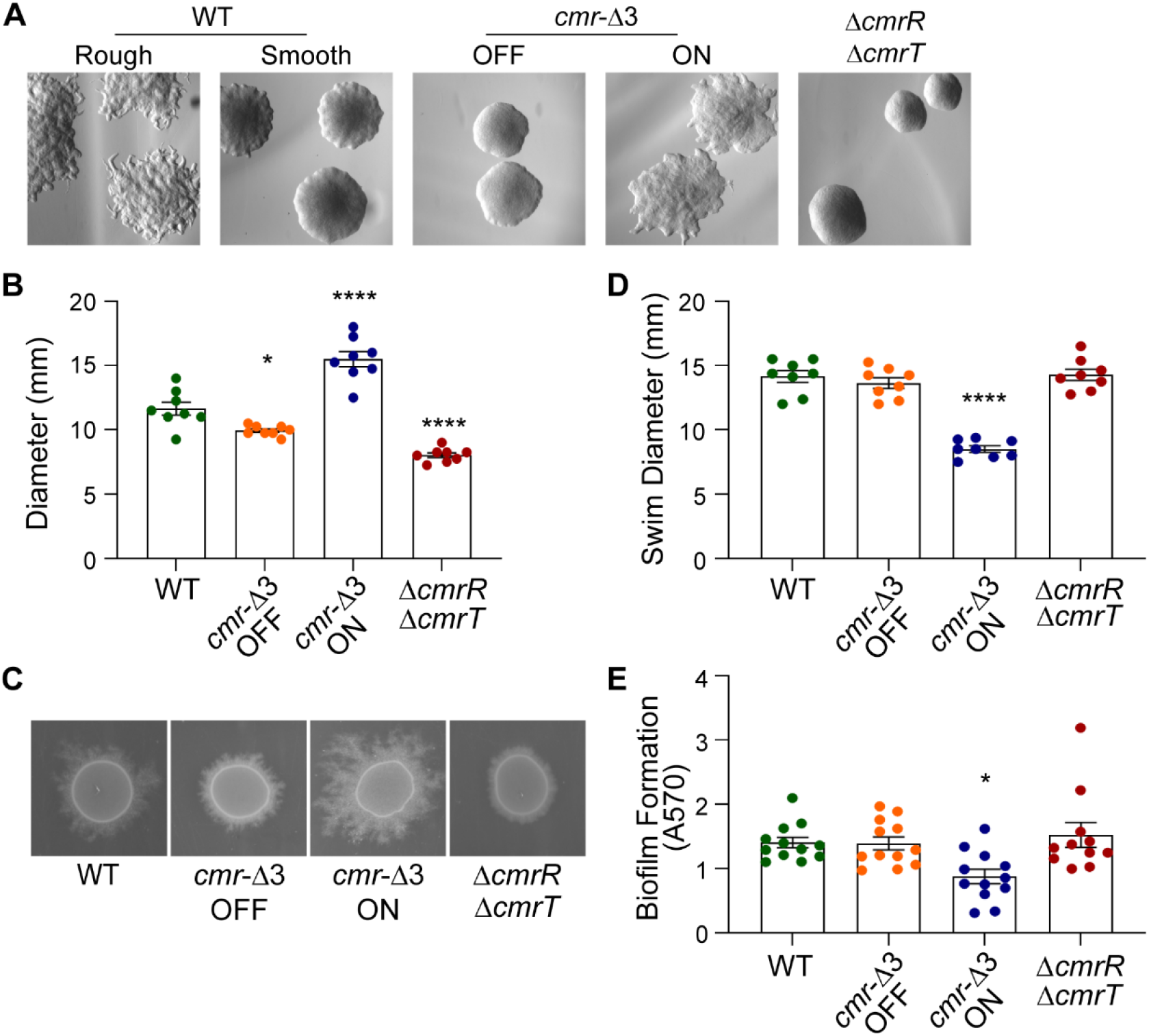
Mutating the RIR of the *cmr* switch results in phenotypically phase-locked strains. (A) Colony morphology of WT rough and smooth isolates, *cmr*-Δ3 OFF, *cmr*-Δ3 ON, and Δ*cmrR*Δ*cmrT*. Shown are representative images after 24 hours growth on BHIS agar. (B) Surface motility of WT, *cmr*-Δ3 OFF, *cmr*-Δ3 ON, and Δ*cmrR*Δ*cmrT* quantified by measuring the diameter of growth after 72 hours. (C) Representative images of surface motility after 72 hours of growth. (D) Swimming motility of WT, *cmr*-Δ3 OFF, *cmr*-Δ3 ON, and Δ*cmrR*Δ*cmrT* mutant quantified by measuring the diameter of growth in 0.5x BHIS 0.3% agar after 48 hours. E) Biofilm formation after 24 hours in BHIS-1% glucose-50 mM sodium phosphate, quantified by crystal violet staining. (B, D, E) Shown are means and standard error, with symbols representing independent samples from at least 2 independent experiments. *p < 0.05, ****p < 0.0001, one-way ANOVA with Dunnett’s multiple comparisons post-test compared to the WT.

### C-di-GMP regulates *cmrRST* expression independently of the *cmr* switch

Upstream of the *cmr* switch is a σ^A^-dependent promoter followed by a c-di-GMP riboswitch sequence, adding another regulatory layer to the transcription of *cmrRST* (33,34). The two regulatory features could act independently with one regulatory element dominant over the other, or they might act coordinately such that both the riboswitch and *cmr* switch must be in the “on” state to allow *cmrRST* transcription. To distinguish between these possibilities, we modulated c-di-GMP levels in the phase-locked *cmr*-Δ3 ON and -OFF strains by overexpressing the diguanylate cyclase *dccA* (pP_tet_::*dccA*) to increase intracellular c-di-GMP or the catalytic EAL domain of the phosphodiesterase *pdcA*, which hydrolyzes c-di-GMP, to reduce c-di-GMP (pP_tet_::EAL) (10,38,39). Increasing c-di-GMP through overexpression of *dccA* resulted in rough colony formation in *cmr*-Δ3 OFF, which normally forms smooth colonies (Figure 3A). Additionally, increasing c-di-GMP enhanced the rough colony appearance of *cmr*-Δ3 ON. In contrast, reduction of c-di-GMP through overexpression of the EAL domain did not visibly affect the morphology of *cmr*-Δ3 ON or -OFF, suggesting that the impact of EAL overexpression on c-di-GMP levels was insufficient to alter expression of *cmrRST* compared to the WT with vector under these conditions. Overexpression of *dccA* led to a significant 2- to 3-fold increase in the abundance of the *cmrS* transcript, which served as a marker for *cmrRST* expression, in both *cmr*-Δ3 ON and -OFF, consistent with the colony morphology phenotypes observed (Figure 3B).

**Figure 3.**
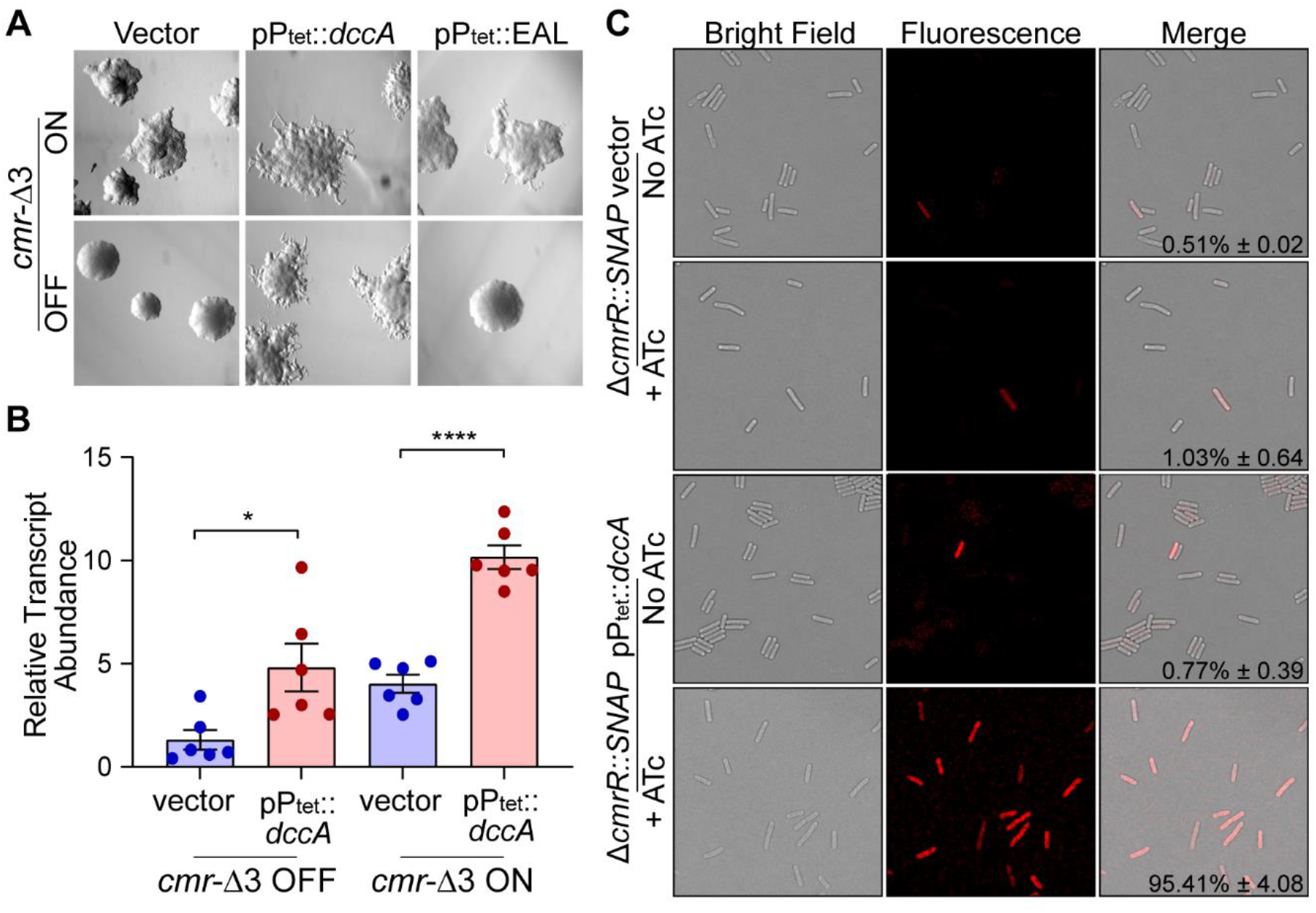
Regulation of *cmrRST* by c-di-GMP occurs independent of *cmr* switch orientation. (A) Colony morphology of *cmr*-Δ3 OFF and *cmr*-Δ3 ON containing either vector, pP_tet_::*dccA*, or pP_tet_::EAL. Strains were grown on BHIS-agar with 20 ng/ml ATc to induce DGC and EAL gene expression. (B) qRT-PCR analysis of *cmrR* transcript abundance in *cmr*-Δ3 OFF and *cmr*-Δ3 ON strains carrying pP_*tet*_::*dccA* or vector control grown in BHIS broth with 20 ng/ml ATc. Data were analyzed using the ΔΔCt method with *rpoC* as the reference gene and normalization to *cmr*-Δ3 OFF with vector. Shown are means with standard error of 6 biological replicates from two independent experiments. *p < 0.05, ****p < 0.0001, one-way ANOVA with Tukey’s multiple comparisons post-test. C) Representative images of Δ*cmrR*::SNAP carrying pP_tet_::*dccA* or vector control was grown in BHIS broth with 20 ng/ml ATc. Samples were imaged at 60x magnification by light and fluorescence microscopy following staining with SNAP-Cell TMR-Star substrate. Percentages of total cells that were fluorescence-positive were determined from at least three fields each from two biological replicates. Shown are means and standard deviations.

These methods assess changes in *cmrRST* expression as a population average. To determine the effects of the c-di-GMP riboswitch and the *cmr* switch on the heterogeneity of *cmrRST* expression among individual bacteria, we used a strain in which *cmrR* was replaced by a codon-optimized *SNAP-*tag gene reporter, Δ*cmrR*::SNAP (25). The pP_*tet*_::*dccA* plasmid and vector control were introduced into this strain. In Δ*cmrR*::SNAP with vector and in the uninduced control, approximately 1% of the population fluoresced (Figure 3C). This result is similar to prior analyses showing that a minority of WT cells express *cmrRST* (25). Induction of *dccA* increased fluorescence-positive cells to > 95%. This result was not due to a shift in *cmr* switch orientation; by qPCR, the orientation of the *cmr* switch was not significantly different between WT overexpressing *dccA* and the vector control (Figure S1). Therefore, the higher percentage of fluorescent cells reflects *cmrRST* expression in response to c-di-GMP. Together, these data indicate that expression of *cmrRST* from the upstream σ^A^-dependent promoter and c-di-GMP riboswitch is not dependent on the orientation of the *cmr* switch. Rather, high c-di-GMP levels increase *cmrRST* expression regardless of *cmr* switch orientation.

### The *cmr* switch contains a promoter in the ON orientation

The best characterized phase variation regulatory mechanisms involve a promoter encoded within the invertible element that drives expression of adjacent genes when in the proper orientation (19). Our data suggest that *cmrRST* transcription originates from multiple sites in a manner dependent on internal c-di-GMP concentrations and the orientation of the *cmr* switch. Therefore, we sought to map the transcriptional start sites (TSS) in strains reflecting these different states. Instead of using *cmr*-Δ3 phase locked strains, which are missing three nucleotides of unknown significance to transcription, we used strains in which the *cmr* switch is locked by a mutation in *recV*, designated *recV cmr*-ON and *recV cmr*-OFF (Figure S2). Additionally, *dccA* was overexpressed in *recV cmr*-OFF to identify any TSS associated with increased c-di-GMP and transcriptional readthrough via the riboswitch. Using 5’ RACE, four total TSS were identified in *recV cmr*-ON, *recV cmr*-OFF, and *recV cmr*-OFF pDccA. In *recV cmr*-OFF with and without *dccA* expression, a TSS designated TSS1 was identified 699 nt upstream of the *cmrR* start codon, located upstream of the c-di-GMP riboswitch (Figure 4A, Figure S3). TSS1 was not detected in *recV cmr*-ON, but this may be due to lower relative abundance of the TSS1 transcript in this strain compared to transcripts from other initiation sites. Another TSS, designated TSS2, was found in *recV cmr*-ON, located 336 nt upstream of *cmrR* and mapping within *cmr* switch in the ON orientation. A third TSS, designated TSS3, was identified 342 nt upstream of the *cmrR* start codon in both *recV cmr*-OFF strains. This position maps TSS3 to within the *cmr* switch, appearing only in transcripts from the *cmr*-OFF strains. In all three strains, a fourth TSS (TSS4) was identified 90 nt upstream from the *cmrR* start codon.

**Figure 4.**
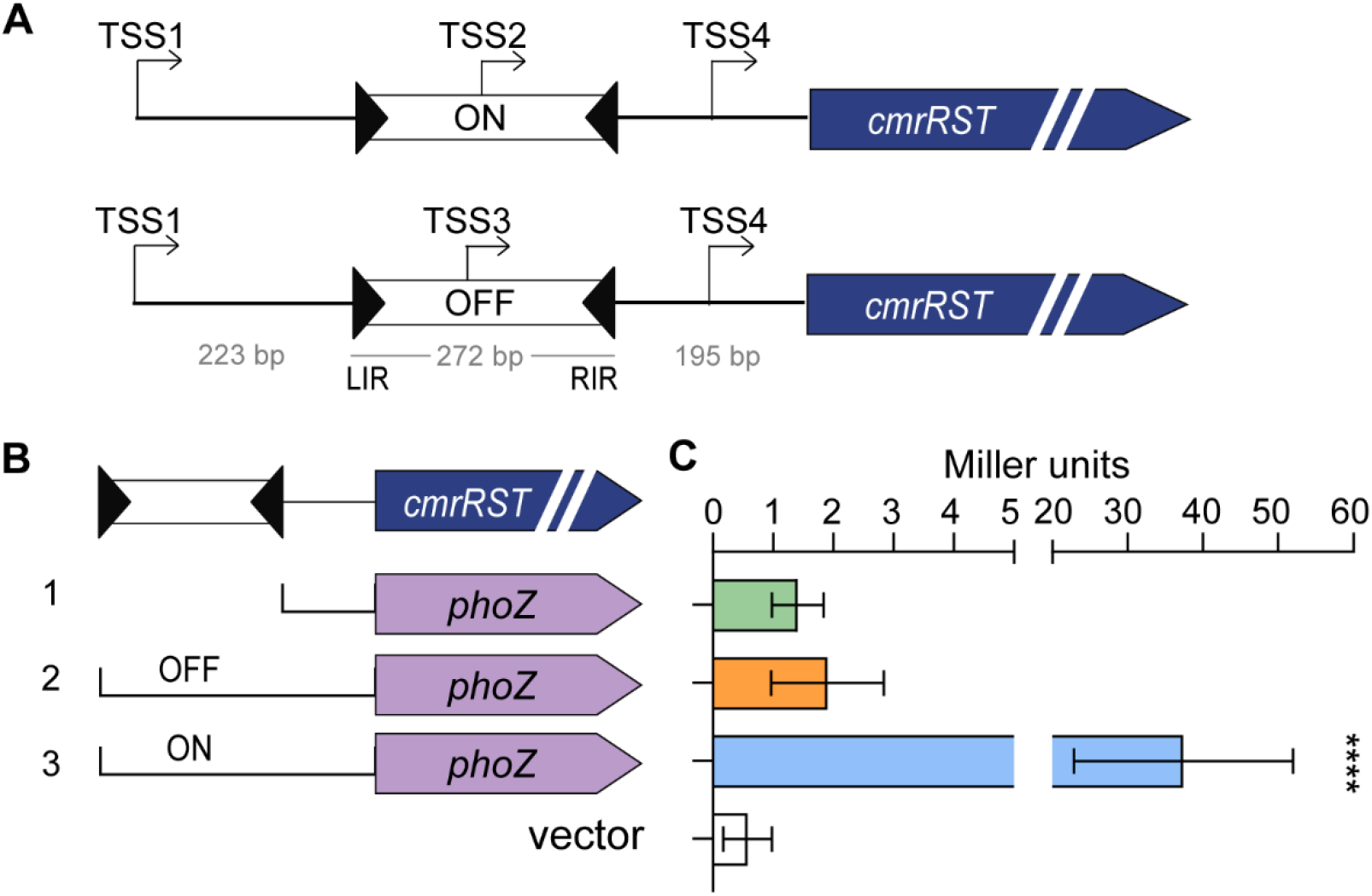
Mapping of transcriptional start sites (TSS) upstream of *cmrRST*. (A) Diagram of TSS identified by 5' RACE using RNA derived from *cmr* ON and *cmr* OFF templates. (B) Alkaline phosphatase reporters used to detect promoter activity. Brackets indicate the regions from the native *cmr* locus that were used to make transcriptional reporters. Construct 1 contains the region between the 3’ end of the right inverted repeat and the 5’ end of *cmrR* coding sequence (pMC123::TSS4-*phoZ*). Construct 2 includes the region in construct 1 plus the *cmr* switch in the OFF orientation (pMC123*::cmr*OFF-*phoZ*). Construct/TSS4 3 is identical to construct 2 but with the *cmr* switch in the ON orientation (pMC123*::cmr*ON/TSS4-*phoZ*). Shown are the means and standard deviations from 12 biological replicates from four independent experiments. ****p < 0.0001, one-way ANOVA with Dunnett’s multiple comparisons test.

The 5’ RACE results confirm an expected TSS upstream of the c-di-GMP riboswitch and suggest that expression of *cmrRST* can originate from multiple additional sites. TSS2 within the *cmr* switch in the ON orientation was identified, however we unexpectedly identified TSS3 within the *cmr* switch in the OFF orientation. To elucidate how phase variable *cmrRST* expression is mediated, we examined the functionality of the newly identified TSS. We created a series of transcriptional reporter fusions of the alkaline phosphatase (AP) gene *phoZ* to regions immediately upstream of *cmrRST* (Figure 4B) (40). Two of the fusions contain the upstream region from the left inverted repeat (LIR) to the start of the *cmrR* coding sequence. These fusions differ in the orientation of the invertible element: ON (pMC123*::cmr*ON/TSS4-*phoZ*) encompasses TSS2, and OFF (pMC123*::cmr*OFF/TSS4-*phoZ*) encompasses TSS3. Both constructs also include TSS4. Additionally, we included a reporter with the region from the 3’ end of the RIR to the start of the *cmrR* coding sequence (pMC123::TSS4-*phoZ*), which contains TSS4, as well as a promoterless control (vector). To prevent inversion of the *cmr* switch present in the reporters once in *C. difficile*, we assayed AP activity in a *recV* mutant lacking the recombinase necessary for inversion (25). *C. difficile* with pMC123::TSS4-*phoZ* (Figure 4B, construct 1) or pMC123*::cmr*OFF/TSS4-*phoZ* (construct 2) showed no significant difference in activity compared to the promoterless control (Figure 4B). In contrast, *C. difficile* with pMC123*::cmr*ON/TSS4-*phoZ* (construct 3) exhibited 20-fold higher AP activity (Figure 4B). These results indicate the presence of a functional promoter when the *cmr* switch is in the ON orientation but not the OFF orientation.

### CmrR positively autoregulates expression of *cmrRST*

Response regulators often have the property of autoregulating their expression (41). To determine whether CmrR and/or CmrT have the ability to transcriptionally regulate *cmrRST*, we overexpressed *cmrR* (pCmrR) and *cmrT* (pCmrT) in *C. difficile* and analyzed *cmrRST* expression by qRT-PCR. Overexpression of *cmrR* resulted in a significant 7-fold increase in transcript abundance of *cmrS* and *cmrT* as compared to the vector control (Figure 5A). In contrast, overexpression of *cmrT* had no effect on *cmrR* or *cmrS* transcript abundance. These results indicate that CmrR, but not CmrT, transcriptionally regulates the *cmrRST* operon.

**Figure 5.**
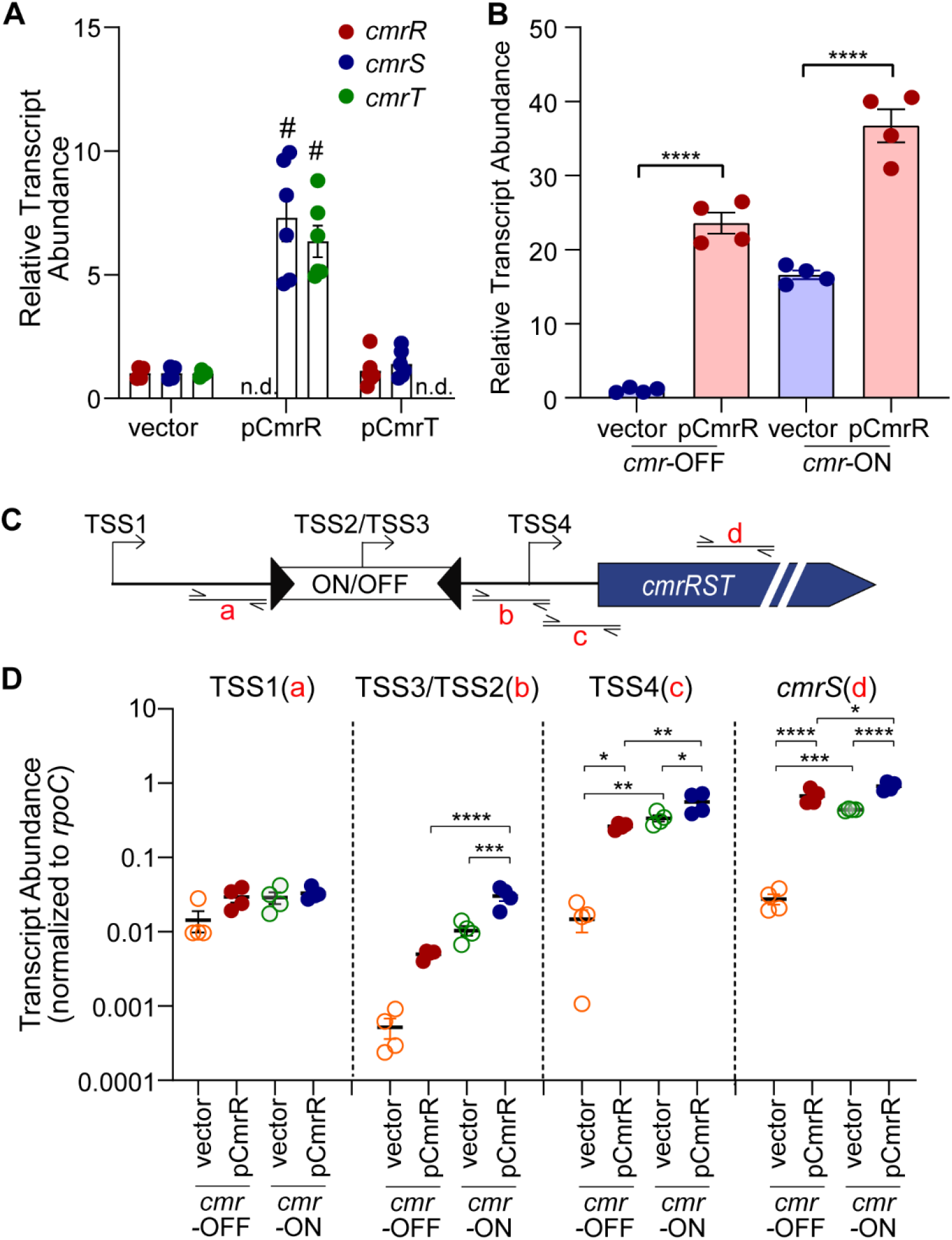
CmrR positively regulates *cmrRST* expression. qRT-PCR measurement of *cmrRST* transcript abundance. Strains were grown with ATc induction (10 ng/mL for vector and pCmrR, 2 ng/mL for pCmrT) prior to RNA isolation. (A) WT with pCmrR, pCmrT, and vector control. n.d., not determined; *cmrR* and *cmrT* were not measured in strains overexpressing those genes. ****p < 0.0001, unpaired t-test. (B) qRT-PCR analysis of *cmrS* transcript abundance of *recV cmr* OFF and *cmr* ON, each with pCmrR or vector control. The *recV cmr*-OFF with vector strain serves as the reference condition for ΔΔCt analysis. Shown are means and standard deviations of 4 biological replicates. (C) Diagram showing regions of transcript measured in (D). (D) qRT-PCR analysis of *cmrRST* transcripts with values normalized to *rpoC*. (B, D) Shown are means and standard error from independent replicates. *p < 0.05, **p < 0.01, ***p < 0.001, ****p < 0.0001, one-way ANOVA with Tukey’s multiple comparisons.

To assess whether CmrR-mediated regulation is dependent on the orientation of the invertible element, we overexpressed *cmrR* from a plasmid in the phase-locked *recV cmr*-ON and *cmr*-OFF mutants. We then used qRT-PCR measuring *cmrS* to assess expression. Overexpression of *cmrR* significantly increased *cmrS* transcript abundance compared to the respective vector control in both *recV cmr*-OFF and *cmr*-ON mutants (Figure 5B). While overexpression of *cmrR* in *recV cmr*-OFF resulted in over a 20-fold increase in *cmrS* transcript abundance relative to vector control, the *cmrS* transcript increased only about 2-fold in *recV cmr-*ON pCmrR relative to vector control.

We considered the possibility that CmrR regulates *cmrRST* expression from one of the previously identified TSS. To address this possibility, we used qRT-PCR to measure the transcript abundance by probing sequences immediately downstream of TSS1, the *cmr* switch containing TSS2 or TSS3, and TSS4, as well as within *cmrS* (Figure 5C). Because transcript abundance differed substantially depending on the region measured, we expressed the data normalized to *rpoC* transcript abundance rather than the ΔΔCt method. Overexpression of *cmrR* did not alter transcript abundance when probing immediately downstream of TSS1, regardless of *cmr* switch orientation, indicating that CmrR-mediated autoregulation occurs downstream of the TSS1 promoter (Figure 5D). In contrast, when probing immediately downstream of the *cmr* switch (TSS2/TSS3), *cmrR* overexpression resulted in 11-fold higher transcript abundance in *recV cmr*-OFF relative to the vector control, though the difference did not reach statistical significance (Figure 5D). Transcript abundance was 23-fold higher in *recV cmr*-ON compared to *recV cmr*-OFF (vector controls, p < 0.01), and *cmrR* overexpression in the *cmr*-ON background led to a further 3-fold increase. Similar changes were observed when probing downstream of TSS4: *cmrR* overexpression increased transcript abundance 31-fold in *recV cmr*-OFF and 60-fold higher in *recV cmr*-ON compared to the *recV cmr*-OFF vector control (Figure 5D). These results suggest that CmrR autoregulates *cmrRST* expression from the promoter corresponding to TSS2, TSS4, or both. Notably, transcript abundance was 29- and 36-fold higher when measuring downstream of TSS4 compared to TSS3 (*cmr*-OFF) and TSS2 (*cmr*-ON), respectively, indicating that the TSS2 promoter is weaker than the TSS4 promoter under these conditions.

### CmrR autoregulation of *cmrRST* expression requires the TSS4 promoter

To further define the promoter autoregulated by CmrR, we generated reporter strains in which different portions of the *cmrRST* regulatory region were transcriptionally fused to a *phoZ* (alkaline phosphatase, AP) reporter gene (40). We first constructed a strain in which *cmrR,* under the control of the P_*tet*_ promoter, was inserted at an ectopic site on the RT1693 (*recV cmr-*OFF) chromosome. This strain allowed ATc-inducible expression of *cmrR* (Figure S4), prevented unintended inversion of the *cmr* switch (Figure S2), and permitted the use of plasmid-borne *phoZ* reporters. Figure 6 shows the combinations of the TSS1/c-di-GMP riboswitch, the *cmr* switch orientation (TSS2/ON or TSS3/OFF), and the region encompassing TSS4.

**Figure 6.**
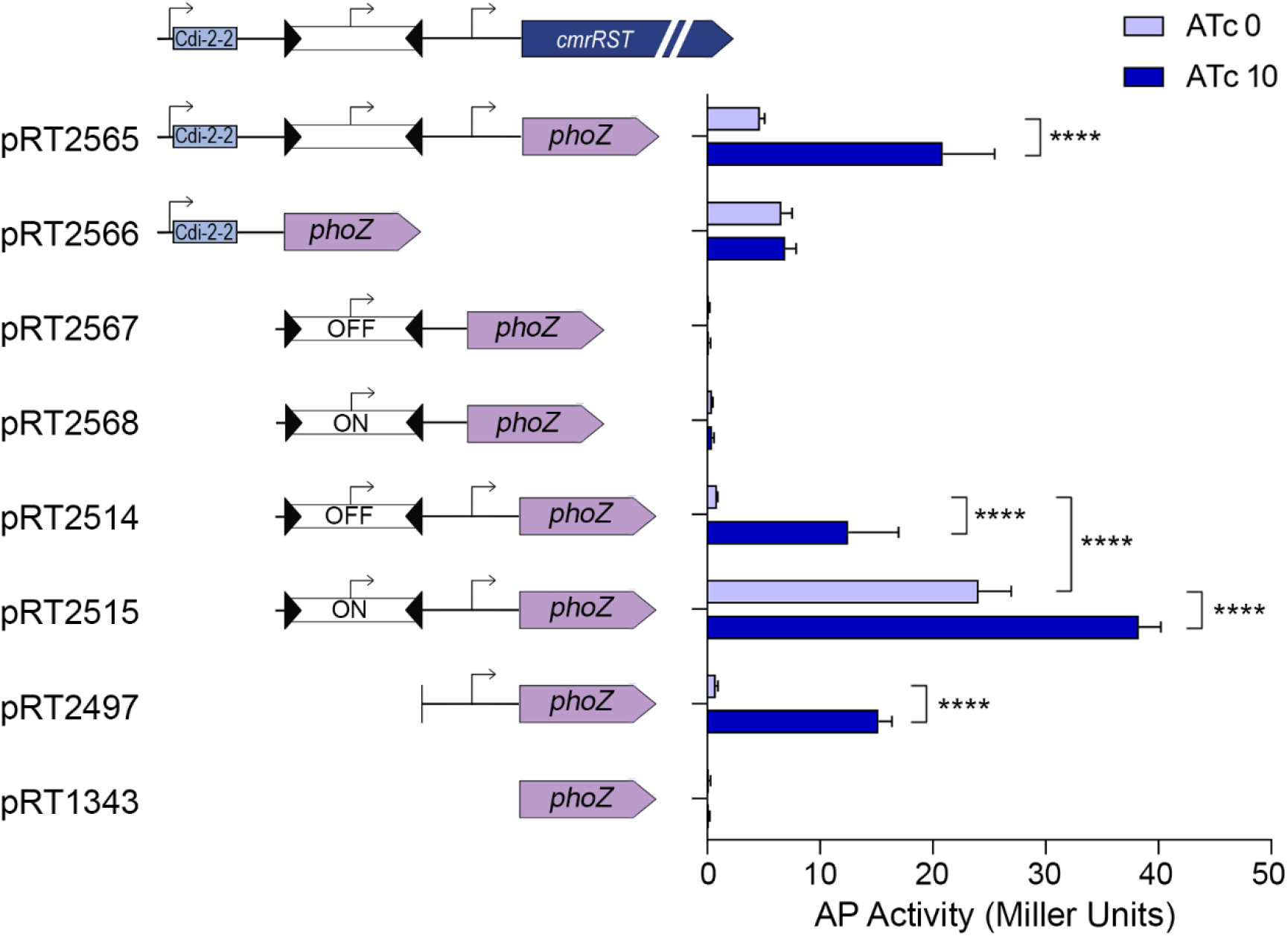
CmrR autoregulation occurs at the TSS4 promoter and is enhanced by the *cmr* ON switch sequence. Alkaline phosphatase reporter activity for regions of the *cmr* 5’ UTR containing one or more putative promoter in *C. difficile* with ATc-inducible Ptet::*cmrR* at an ectopic site. (Left) Diagrams of the plasmid-borne reporter fusions carried by the strain. (Right) AP reporter activity after growth with or without 10 ng/mL ATc to induce cmrR expression (Figure S4). Shown are means and standard deviations from three independent experiments. ****p < 0.0001, two-way ANOVA and Sidak’s post-test.

The promoter corresponding to TSS1, located upstream the c-di-GMP riboswitch, displayed similar transcriptional activity regardless of ATc (pRT2566), indicating that CmrR does not autoregulate expression at this site. This result is consistent with the qRT-PCR analysis of the native *cmrRST* site (Figure 5), and it further suggests that *cmrR* overexpression does not impact global intracellular c-di-GMP levels. The promoter corresponding to TSS3 (pRT2567), present in the *cmr* OFF switch orientation, yielded no detectable AP activity when isolated from upstream (TSS1) and downstream (TSS4) sequences and was not affected by overexpression of *cmrR*. Adding the downstream sequence to the *cmr* OFF switch region (TSS3 + TSS4, pRT2514) resulted in no AP activity in the absence of ATc but led to 14-fold increased AP activity when *cmrR* was overexpressed.

Interestingly, the construct containing only the *cmr* ON switch sequence (TSS2, pRT2568) also lacked transcriptional activity with and without *cmrR* overexpression. The addition of the downstream sequence (TSS2 + TSS4, pRT2515) yielded baseline AP activity 52-fold higher than the TSS2-only reporter and also conferred CmrR-mediated autoactivation. Finally, the reporter containing the region downstream of the *cmr* switch alone (TSS4, pRT2497) did not show AP activity in the absence of ATc but did exhibit 19-fold activation by *cmrR* overexpression. The baseline AP activity and degree of activation by CmrR was comparable to that from pRT2514 (TSS3 + TSS4), confirming that there is no active promoter in the *cmr* switch when in the OFF orientation under these conditions. Overall, only the constructs with the TSS4 region resulted in increased activity upon *cmrR* overexpression, indicating that the TSS4 promoter is required for CmrR-mediated autoregulation of *cmrRST*. Further, the highest transcriptional activity in the absence of *cmrR* overexpression was seen in the *cmr* ON construct in which TSS2 and TSS4 are both present, suggesting additive effects of the promoters or the disruption of an important sequence in the constructs separating the two elements.

### *cmrRST* and its upstream regulatory elements are well conserved in *C. difficile*

*C. difficile* strains from multiple ribotypes form rough and smooth colonies associated with phase variable *cmrRST* expression (32). To examine the potential scope of CmrRST phase variation and function in *C. difficile*, we used NCBI BLAST to determine the extent of conservation of *cmrRST* and its upstream regulatory region using R20291 as the reference sequence. The analysis examined the 895 bases 5’ of *cmrR*, which contains a c-di-GMP riboswitch and the *cmr* switch (Figure 1A), and the *cmrRST* coding sequences. Among 71 *C. difficile* whole genome sequences available in NCBI (taxonomic ID 1496), *C. difficile* strains shared high sequence identity to the reference for both the riboswitch (98.9% ± 1.2) and the *cmr* invertible element (98.1% ± 1.4) (Figure S5). This similarity is comparable to the average percent sequence identity for the *cmrRST* operon (98.7% ± 1.1). These results indicate that the c-di-GMP riboswitch and invertible element upstream of *cmrRST* are highly conserved across many *C. difficile* strains from divergent ribotypes and underscores the importance of this operon and its regulation to *C. difficile* physiology.

## Discussion

In this study, we demonstrate that multiple regulatory mechanisms control the transcription of *cmrRST*. Our results support a model in which, under basal c-di-GMP conditions, the orientation of the *cmr* switch determines the expression level of *cmrRST*. This phase variation mechanism involves the reversible inversion of a promoter within the *cmr* switch sequence. Increasing intracellular c-di-GMP augments *cmrRST* expression independent of *cmr* switch orientation. Our data also show that expression of *cmrRST* is also subject to autoregulation by CmrR, which occurs at an additional promoter downstream of the *cmr* switch and is enhanced by the orientation of the *cmr* switch. Therefore, multiple environmental signals may impact *cmrRST* expression through c-di-GMP signaling, phase variation, and the activation of the response regulator CmrR. CmrRST has important roles in *C. difficile* cell and colony morphology, motility, biofilm formation, and virulence, suggesting multiple contexts in which distinct environmental stimuli and selective pressures must be integrated to appropriately control *cmrRST* expression at the single-cell and population levels. This work represents the first account of a unique intersection of regulatory mechanisms controlling the expression of a signal transduction system that broadly impacts *C. difficile* physiology and disease development.

To disentangle the roles of c-di-GMP and phase variation in *cmrRST* transcription, we phase locked the *cmr* switch by deleting three nucleotides in the RIR, one of which was identified as the precise site of recombination during inversion of the *cmr* switch (25). Phenotypic analysis of phase locked R20291 *cmr*- Δ3 ON and *cmr*-Δ3 OFF mutants showed that they behave similarly to WT rough and smooth populations, respectively. Expression of *cmrRST* in the locked ON mutant was equivalent to that of WT rough isolates, while expression in the *cmr*-Δ3 OFF mutant was similar to that of WT smooth isolates. Consistent with these results, *cmr*-Δ3 ON yielded rough colonies, increased surface motility and decreased swimming motility and biofilm formation as compared to WT smooth isolates, the *cmr*-OFF mutant, and the Δ*cmrR*Δ*cmrT* mutant. These data indicate that orientation of the *cmr* switch controls expression of *cmrRST* and further confirms the role of CmrRST in these phenotypes. Interestingly, while *cmr*-Δ3 ON forms rough colonies, they are not identical to those formed in WT populations, and there is still some heterogeneity of colony morphology. This observation may reflect differences in activity of CmrR or CmrT or that other factors contribute to colony morphology. Other work has suggested heterogeneity in c-di-GMP levels among individual *C. difficile* cells (21,24), so c-di-GMP may also result in colony morphology differences via modulation of *cmrRST* expression.

By artificially manipulating intracellular c-di-GMP levels in the *cmr* locked mutants, we found that c-di-GMP promotes *cmrRST* expression regardless of the orientation of the *cmr* switch. These findings are consistent with observed phenotypes; the *cmr*-Δ3 OFF mutant exhibited surface motility intermediate between that of WT and Δ*cmrR*Δ*cmrT* mutant. In the *cmr*-Δ3 OFF mutant, *cmrRST* could still be expressed from the TSS1 promoter upstream of the invertible element if c-di-GMP levels are sufficiently high. Intracellular c-di-GMP levels have been shown to increase in *C. difficile* with growth on a surface (39), which is consistent with c-di-GMP serving as a signal to enhance CmrRST-mediated surface motility regardless of *cmr* switch orientation.

Multiple new TSS, in addition to the previously identified σ^A^-dependent promoter corresponding to TSS1 upstream of the c-di-GMP riboswitch (33,34), were identified upstream of *cmrRST* using 5’ RACE: TSS2 was identified within the *cmr* switch in *cmr*-ON *C. difficile*, TSS3 was detected within the *cmr* switch in *cmr*-OFF, and TSS4 was found between the *cmr* switch and the translational start of *cmrR* in both strains. TSS3 may have been a spurious result, as the associated putative promoter lacked detectable transcriptional activity, and no −10/−35 sites similar to σ^70^ consensus sequences were identifiable upstream of TSS3. The TSS2 region, when isolated from the TSS1 and TSS4 regions, also lacked detectable transcriptional activity using the AP assay, and transcript abundance measured by qRT-PCR was similarly low for the region immediately downstream of TSS2. TSS2 may nonetheless indicate a weak promoter, supported by the presence of −10 and −35 sites upstream of TSS2 (Figure S3) and its responsiveness to CmrR. The level of TSS2 promoter activity, even with CmrR activation, may have been below the detectable limit using the *phoZ* reporter. A longer region containing both TSS2 and TSS4 showed maximal reporter activity, indicating that both sequences must be present to allow optimal transcription initiation. This effect was not observed for the equivalent TSS3 + TSS4 region, suggesting that the ON *cmr* switch sequence contains information that enhances expression. Further, only the reporters containing TSS4 exhibited CmrR-mediated autoactivation. The TSS4 region alone or with the upstream TSS3 *cmr* OFF sequence showed equivalent increases in activity upon over-expression of *cmrR*. Together these results support a model (Figure 7) in which the TSS1 promoter (P1) acts independently of the downstream promoters and yields an mRNA with a c-di-GMP riboswitch that enhances transcription readthrough when the c-di-GMP ligand is bound. The *cmr* switch in the ON state contains a weak promoter (P2) corresponding to TSS2 that can be activated by CmrR; the inverse orientation of the *cmr* switch lacks a promoter. An additional promoter (P4) downstream of the *cmr* switch is also activated by CmrR and strongly contributes to autoactivation of *cmrRST*, particularly when the *cmr* switch is in the ON orientation. These regulatory elements are poised to modulate *cmrRST* expression in response to distinct environmental stimuli. Interestingly, CmrT does not regulate the expression of *cmrRST*, suggesting that CmrR and CmrT have different DNA specificities and, accordingly, distinct regulons. Future work will identify the genes directly and indirectly regulated by CmrR, CmrT, or both as well as determine their consensus DNA binding sites, which will elucidate the functions of these co-expressed response regulators.

**Figure 7.**
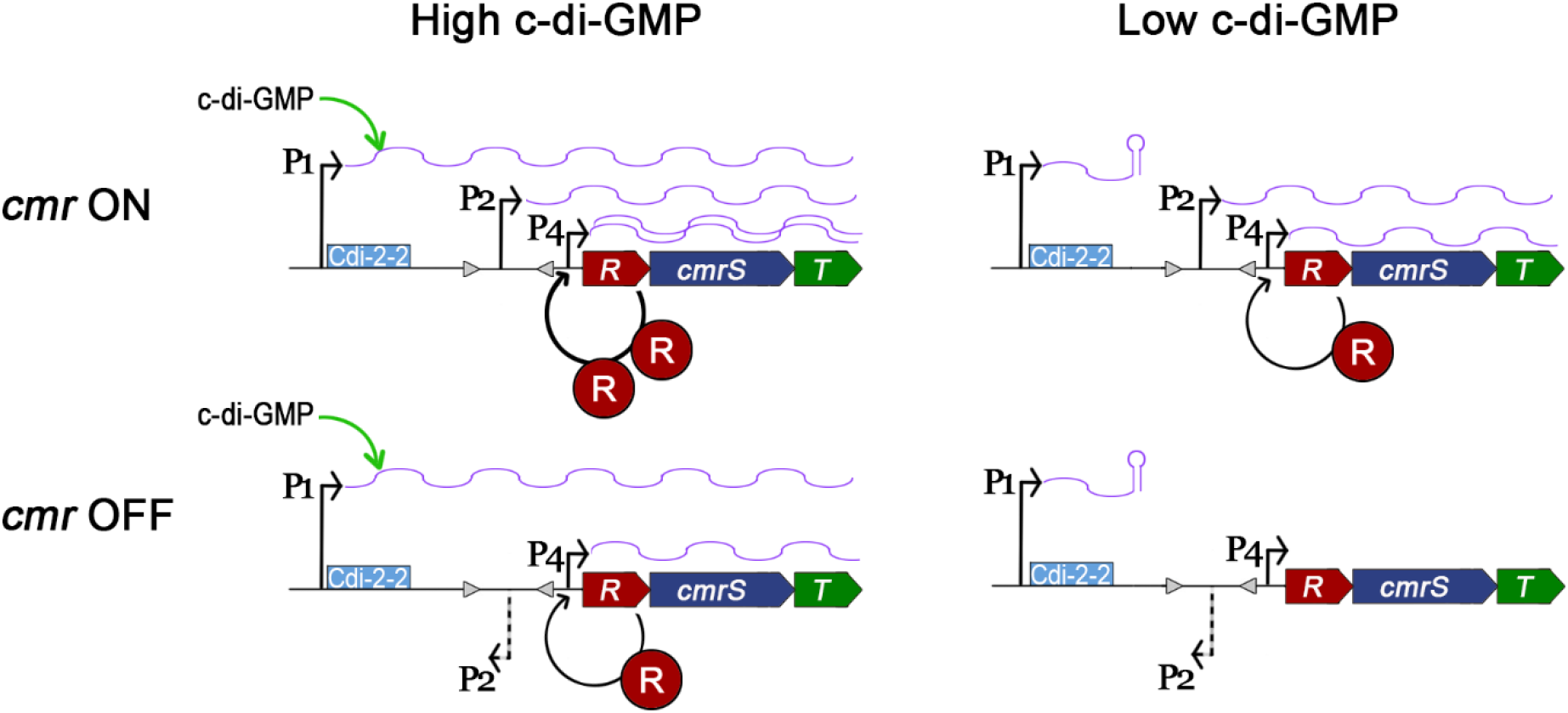
Model of multilevel regulation of *cmrRST* transcription. Three promoters control the expression of *cmrRST*. Promoter P1 corresponding to TSS1 generates an mRNA with a c-di-GMP riboswitch that is permissive for transcriptional readthrough when c-di-GMP is bound. The *cmr* switch contains a weak promoter (P2) corresponding the TSS2 when in the ON orientation. An additional promoter, P4, lies between the *cmr* switch and the *cmrRST* coding sequences. Transcription from P4 is positively regulated by CmrR. P1 and P2 independently regulate *cmrRST* transcription, while P4 activity is enhanced when the *cmr* switch is in the ON orientation.

In other bacteria, the reversible inversion of a promoter via site-specific DNA recombination is common mechanism of phase variable expression of adjacent genes (15,42). The *cmr* switch is the first instance of this mechanism of regulation in *C. difficile*, though the inversion of the TSS2 promoter may be less impactful to *cmrRST* transcription than other features of the *cmr* ON sequence combined with the downstream TSS4 promoter. Notably, the region upstream of the *flgB* operon has a similar arrangement to that of *cmrRST*, with a c-di-GMP riboswitch preceding the switch that undergoes site-specific recombination. the flagellar switch that mediates phase variation (21,27,34). However, the riboswitch upstream of *flgB* negatively regulates transcriptional readthrough, so binding of c-di-GMP by the riboswitch leads to transcription termination precluding synthesis of longer transcripts containing the flagellar switch regardless of its orientation. Thus, phase variation and c-di-GMP regulation of flagellar gene expression are linked, in contrast with the *cmrRST* system in which c-di-GMP and the *cmr* switch independently modulate expression.

In summary, this work demonstrates that *cmrRST* expression is subject to multilayered regulation with multiple potential inputs from environmental signals. The complexity of this regulatory network suggests that *cmrRST* expression, and therefore its transcriptional targets, requires careful control. Further work that defines the signals which promote *cmrRST* expression will provide important insights into the role of this TCS in *C. difficile* physiology and pathogenesis.

## MATERIALS AND METHODS

### Growth and maintenance of bacterial strains

Table S1 lists strains and plasmids used in this study. *C. difficile* R20291 and derivative strains were maintained in an anaerobic environment of 90% N_2_, 5% CO_2_, and 5% H_2_. *C. difficile* strains were grown statically at 37 °C in BHI, BHIS (37 g/L Bacto brain heart infusion, 5 g/L yeast extract), or in Tryptone Yeast (TY; 30 g/L Bacto tryptone, 20 g/L yeast extract, 1 g/L thioglycolate) medium as indicated. *E. coli* strains were grown in Luria Bertani medium at 37 °C with aeration. Antibiotics were used where indicated at the following concentrations: chloramphenicol (Cm), 10 μg/mL; thiamphenicol (Tm), 10 μg/mL; kanamycin (Kan), 100 μg/mL; ampicillin (Amp), 100 μg/mL.

### Construction of bacterial strains and plasmids

Table S2 lists primers used in this study. Detailed methods for the generation of plasmids and strains used in this study are provided in Supplemental Methods. Strain and plasmid information is listed in Table S1.

### Quantitative reverse-transcriptase PCR

For analysis of *cmrR* and *cmrT* expression in *C. difficile* R20291 (WT), *cmr*-Δ3 ON and *cmr*-Δ3 OFF, these strains were grown overnight (16 h) in TY medium, and 5 μL were spotted on BHIS-agar. After 24 hours, growth was collected, suspended in 1:1 ethanol:acetone, and stored at −80 °C for subsequent RNA isolation. For analysis of transcript abundance in R20291 strains carrying pCmrR, pCmrT, or vector, these strains were grown overnight in TY-Tm medium. Cultures were diluted 1:30 in BHIS-Tm broth. After two hours growth, ATc was added to induce gene expression (WT with vector or pCmrR, 10 ng/mL; WT with pCmrT, 2 ng/mL). Samples were collected at mid-exponential phase and saved in 1:1 ethanol:acetone at −80 °C.

RNA was extracted as described previously (43) and treated with TURBO DNA-free™ Kit (Life Technologies) to remove contaminating genomic DNA. cDNA was synthesized using the High-Capacity cDNA Reverse Transcription Kit (Applied Biosystems) using the manufacturer’s protocols (22,38). Real-time PCR was performed using SensiFAST SYBR & Fluorescein Kit (Bioline) as previously described (25,32). Data were analyzed using *rpoC* as the reference gene. Primers are as follows: *rpoC*, R850/R851; *cmrR*, R2298/R2299; *cmrT*, R2537/R2538; *cmrS*, R2539/R2540; TSS1, R2745/R2746; TSS2/3, R2751/R2804; TSS4, R2803/R2716.

### Orientation specific PCR

Orientation specific PCR was done as previously described (21,32). Colonies grown on BHIS-agar were boiled to produce lysate for PCR using primers R2270/R2271 to detect the ON orientation (241 bp product) and R2271/R2272 to detect the OFF orientation (140 bp product). PCR products were separated on a TAE-1.5% agarose gel.

### Quantification of switch orientation by quantitative PCR

R20291 *cmrR*::SNAP pP_tet_::*dccA* (RT2500) and vector control (RT2501) were grown overnight in TY-Tm broth then diluted 1:30 in BHIS-Tm containing 20 ng/μL to induce *dccA* expression. After growth to mid-exponential phase, genomic DNA was purified by phenol:chloroform:isopropanol extraction and ethanol precipitation. qPCR was performed as previously described with 100 ng of DNA per 20 μL reaction and 100 nM primers (32). Data were analyzed using the ΔΔCt method, with *rpoA* as the reference gene and the indicated control condition. Primers are as follows: ON orientation, R2270/R2271; OFF orientation, R2271/R2272; and *rpoA,* R2273/R2274.

### Microscopy

To image whole colonies, strains were grown on BHIS-Tm agar for 24 hours. For *cmr*-Δ3 ON/OFF with pP_tet_::*dccA*, pP_tet_::EAL, or vector, ATc (20 ng/mL) was included in the agar. Colonies were imaged using a Keyence BZ-X810 microscope or Zeiss Stereo Discovery V8 dissecting microscope with a glass stage.

Fluorescence microscopy of SNAP-labelled strains was performed as previously described (25). R20291 *cmrR*::SNAP pP_tet_::*dccA* or vector control were grown overnight in TY-Tm broth. Cultures were diluted 1:30 in BHIS-Tm. After two hours of growth, ATc (20 ng/mL) was added for induction. Mid-exponential phase samples were pelleted, washed, and incubated with SNAP-Cell TMR-Star (New England Biolabs) at 37 °C for 30 minutes. Samples were further washed and mounted on 1% agarose pads. Imaging was done using a Keyence BZ-X810 microscope with an Olympus 100X PlanFluor objective. Bright field and visualization of red fluorescence with a Chroma filter (ET545/30x, EM620/60m) was done using the same image capture settings for all samples. Cells were counted on at least three fields from two biological replicates for each strain and condition.

### Phenotype assays

Motility and biofilm assays were done as previously described (32,38); further details are provided in Supplemental Methods.

### Alkaline phosphatase assays

Strains carrying *phoZ* reporters were grown in BHIS medium to OD_600_ ~ 1.0, then 1.5 mL of culture was collected, pelleted and frozen at −20 °C. Samples were thawed, and AP activity using the substrate *p*-nitrophenyl phosphate was measured as previously described (21,40).

### 5’ rapid amplification of cDNA ends (RACE)

R20291 *recV cmr-*OFF (RT1693), *recV cmr-*OFF pDccA (RT2184), and *recV cmr-*ON (RT2520) were grown in BHIS to mid-exponential phase. RT2184 was grown with Tm and induced with 1 μg/mL nisin (induces *dccA* expression from the P*cpr* promoter). Samples were collected and frozen in 1:1 ethanol:acetone. RNA was extracted and purified using the RNeasy Mini Kit (Qiagen). cDNA was synthesized with the 5' RACE System for Rapid Amplification of cDNA Ends kit (Thermo Fisher) using the manufacturer’s protocol. Briefly, cDNA was synthesized using primer R2792 and SuperScript II reverse transcriptase. The samples were treated with RNase Mix then column purified and treated with TdT. The products were then PCR amplified with the provided anchor primer and primer R2793, then subjected to Sanger sequencing.

## Acknowledgements

The funders had no role in study design, data collection and analysis, decision to publish, or preparation of the manuscript.

## SUPPLEMENTAL FIGURES

**Figure S1. C-di-GMP does not cause inversion of the *cmr* switch.** Δ*cmrR*::SNAP carrying pP_tet_::*dccA* or vector control was grown to mid-exponential phase in BHIS broth with or without 20 ng/ml ATc. gDNA was collected for qPCR analysis of *cmr* switch orientation. Data are expressed as the percent OFF orientation. Shown are the means and standard deviations of six biological replicates from two independent experiments. No significant differences, two-way ANOVA with Tukey’s multiple comparisons.

**Figure S2. The *cmr* switch is phase locked in *recV*-deficient strains.** Orientation-specific PCR to detect each orientation of the *cmr* switch in WT, *recV cmr-*OFF, and *recV cmr-*ON.

**Figure S3. Identification and mapping of multiple transcriptional start sites upstream of *cmrRST***. Map of TSS identified by 5’ RACE in *recV cmr*-ON, *recV cmr*-OFF and *recV cmr*-OFF pDccA. Gray highlight indicates the Cdi-2-2 riboswitch sequence. Red and blue text denote the OFF and ON orientations of the *cmr* switch, respectively. Bold text indicates the inverted repeats. Green highlights mark TSS. Underlined text indicates putative −10/−35 sites. Yellow highlight denotes the *cmrR* start codon.

**Figure S4. Inducible expression of *cmrR* from an ectopic chromosomal site.** R20291 *recV* CDR2492::P_*tet*_::*cmrR* was grown to mid-exponential phase in BHIS medium with and without 10 μg/mL ATc to induce *cmrR* expression. qRT-PCR was used to measure *cmrR* transcript abundance using 3 different primer sets to distinguish *cmrR* expression from the native site (primers R3111/R3112), from the ectopic site (primers R3113/R3112), and total *cmrR* (primers R2298/R2299). Data are expressed as relative transcript abundance in the ATc condition compared to that without ATc, with normalization to *rpoC* as the reference strain. Induction with ATc resulted in a 9.0-fold increase in *cmrR* mRNA from the ectopic site and a 2.6-fold increase from the native *cmrRST* locus, with a cumulative 12.7-fold increase in *cmrR* transcript. Shown are means and standard error. * p < 0.05, *** p < 0.001, **** p < 0.0001 by unpaired t-test.

**Figure S5. *cmrRST* and upstream regulatory sequences are highly conserved across *C. difficile* strains and ribotypes.** Heat map showing percent sequence identity of 71 sequenced *C. difficile* strains in the NCBI database compared to the R20291 reference sequence (NCBI accession number FN545816.1). If the strain has been ribotyped, the ribotype is indicated on the left. For the *cmr* switch, both orientations were used as the reference and the highest percent sequence identity is shown. The following nucleotides were used as the reference sequence from R20291: riboswitch, 3,736,512-3,736,863; *cmr* switch, 3,735,969-3,736,513; *cmrRST*, 3,732,474-3,735,968.

## Notes

### Competing Interest Statement

The authors have declared no competing interest.

### Summary of Updates

Supplemental Files have been added.

