## Supplemental Methods for "Multiple regulatory mechanisms control the production of CmrRST, an atypical signal transduction system in *Clostridioides difficile*"

### Construction of bacterial strains and plasmids

*C. difficile* R20291 genome sequence (NCBI Accession No. FN545816) was used in the design of all plasmids and strains. To construct $pP_{tet}$-DccA (pRT1587) and $pP_{tet}$-EAL (pRT2444), *dccA* and the catalytic EAL domain of *pdcA* (1) were PCR-amplified from R20291 genomic DNA (*dccA*, primers R1907/R1908; EAL, primers R2689/R2690). Purified PCR products were digested with SacI and BamHI (New England Biolabs) and ligated into a similarly digested pRPF185 (2).

To construct pMC123::TSS4-*phoZ* (pRT2497), pMC123::*cmr*OFF/TSS4-*phoZ* (pRT2514), pMC123::*cmr*ON/TSS4-*phoZ* (pRT2515), pMC123::5'UTR*cmr*OFF-*phoZ* (pRT2565), pMC123::TSS1-*phoZ* (pRT2566), pMC123*::cmr*OFF-*phoZ* (pRT2567), and pMC123*::cmr*ON-*phoZ* (pRT2568), inserts consisting of portions of the 5' UTR sequence were PCR amplified from R20291 genomic DNA using the following primers: pRT2497, R2754/R2342 to amplify the sequence between the right inverted repeat (RIR) of the *cmr* switch and *cmrR* start codon; pRT2514 and pRT2515, R2337/R2342 to amplify the sequence upstream the left inverted repeat (LIR) of the *cmr* switch and the *cmrR* start codon; pRT2565, R2857/R2342 to amplify the entire 5' UTR; pRT2566, R2857/R2858 to amplify TSS1 and the c-di-GMP riboswitch; pRT2567 and pRT2568, R2859/R2860 to amplify the *cmr* switch. PCR products were digested with EcoRI and BamHI (New England Biolabs) and ligated into similarly digested pMC123*::phoZ* (pRT1343). Ligations were transformed into DH5α, and Cm-resistant colonies were screened by PCR with the above primers. Plasmids were confirmed by PCR and sequencing; sequencing was used to identify clones with the *cmr*-OFF or -ON sequence orientation for pRT2514, pRT2567, pRT2565 (*cmr*-OFF), and pRT2515, pRT2568 (*cmr*-ON). Plasmids were transferred to *C. difficile* R20291 by conjugation using *E. coli* HB101(pRK24) as previously described (3,4).

R20291 Δ*cmrR*Δ*cmrT* (RT2296) was made by creating a markerless deletion of *cmrT* in R20291 Δ*cmrR* (RT2256) as previously described (5). Regions up- and downstream of *cmrT* were amplified by PCR (upstream, primers OS268/OS269; downstream, primers OS271/OS288) and joined by Gibson Assembly (New England Biolabs) into PmeI-linearized pMTL-SC7215 (6). The plasmid was introduced into R20291 Δ*cmrR* by conjugation with *E. coli* HB101(pRK24). Mutants were identified by PCR and verified by PCR and sequencing (5,6).

R20291 *recV cmr*-ON (RT2520) was derived from R20291 *recV cmr*-OFF (RT1697) by ectopically expressing *recV* (pRecV) to stimulate *cmr* switch inversion, then screening resulting isolates for the ON orientation of the *cmr* switch. Specifically, RT1697 was grown in the presence of 20 ng/mL ATc to induce *recV* expression, then plated on BHIS without antibiotic selection to allow loss of pRecV. Tm-sensitive colonies, indicating loss of the plasmid pRecV, were identified by replica plating, and *cmr* ON isolates were identified by PCR with orientation-specific primers R2270/R2271.

The *cmr* switch (Cdi6) was locked either in the OFF or the ON configuration in *C. difficile* R20291 using a modified version of the previously described *codA*-based allelic exchange method (6). Primers were designed with overlapping homology to delete three nucleotides in the RIR at position 3,736,176 – 3,736,178 inclusively. To create the strain with the *cmr* switch locked in the OFF configuration (*cmr*-Δ3 OFF), primer pairs OS383/OS384 and OS385/OS386 were used to PCR-amplify flanking homology regions using R20291 *recV cmr*-OFF genomic DNA as the template. The homology arms were then cloned by Gibson Assembly (New England BioLabs) into PmeI-linearized pMTL-SC7215 vector in which the *codA* negative-selection marker was replaced by the holin-endolysin module from a prophage in the *C. difficile* CD630 strain (locus tags CD630_28941 and CD630_28940, NCBI accession number CP010905.2) under the control of an anhydrotetracycline (ATc)-inducible promoter from the pRPF185 plasmid (2). The resulting plasmid was then conjugated into *C. difficile* R20291 strain as described previously. The clones were selected on BHI agar containing 15 µg/mL Tm and 12 µg/mL norfloxacin. Single-crossover integrant clones were purified by three rounds of streaking onto BHI agar with 15 µg/mL Tm and 12 µg/mL norfloxacin followed by double-crossover selection on BHI containing 100 ng/mL ATc. Colonies were screened for the desired mutation with primers OS387/OS137. Genomic DNA from PCR-positive clones was extracted, and the *cmr* switch region was PCR-amplified using primers OS109/OS111 following by Sanger-sequencing of the amplicon for verification. The same strategy was used to lock the *cmr* switch in the ON configuration (strain *cmr*-Δ3 ON) using primer pairs OS388/OS383 and OS389/OS386 to PCR-amplify homology arms from R20291 *recV cmr*-ON genomic DNA template and OS390/OS391 for PCR-screening of presumed double-crossover clones.

R20291 *recV* $P_{tet::}$*cmrR* (RT2826) was generated by inserting *cmrR* under the control of an ATc-inducible promoter on the chromosome of the *recV* mutant strain (RT1693) (7). The $P_{tet}$::*cmrR* fusion was inserted between CDR20291_2492 and CDR20291_2493, convergently encoded neighboring genes with ~700 bp between them using the pMSR0 allelic exchange vector (8). The $P_{tet}$::*cmrR* fusion was PCR amplified from pCmrR (pRT2073) (5) using primers R2918/R2919. The upstream homology arm (overlapping CDR20291_2492) was amplified using primers R2914/R2915; the downstream homology arm (overlapping CDR20291_2493) was amplified using primers R2916/R2917. The primers used included overlapping homology to allow Gibson Assembly (New England BioLabs) of the products with pMSR0 digested with BamHI-HF at 40°C for 1 hour. The Gibson Assembly product was transformed into *E. coli* DH5α, and Cm-resistant colonies were confirmed via PCR and sequencing with primers R2743/R2744 and R2918/R2919. The confirmed plasmid (pRT2825) was introduced into *C. difficile* R20291 *recV cmr*-OFF (RT1693) via conjugation as described above. Single and double-crossovers were done as described (8). The resulting *C. difficile* $P_{tet}$::*cmrR* strain (RT2826) was confirmed by PCR with primers R2987/R2988 and by qRT-PCR to ensure ATc-inducible *cmrR* expression (Figure S4). Plasmids containing *phoZ* reporter fusions (pRT1343, pRT2497, pRT2514, pRT2515, pRT2565, pRT2566, pRT2567, and pRT2568) were then conjugated into the $P_{tet}$::*cmrR* strain via *E. coli* HB101(pRK24) as described above.

**Phenotype assays**

Motility assays were done as previously described (5,9). For surface motility, 5 µL of overnight culture was spotted onto BHIS-1% glucose-1.8% agar. For swimming motility, 1 µL was inoculated into 0.5x BHIS-0.3% agar. For strains carrying plasmids, Tm (10 µg/mL) and ATc induction was included (10 ng/mL for vector and pCmrR; 2 ng/mL for pCmrT). After 48 hours (swimming) or 72 hours (surface), the diameters of the motile spots were measured at the widest point and perpendicular to the first measurement.

These measurements were averaged to reflect overall diameter. Surface motility was imaged after 72 hours using a Syngene G:Box imager.

Biofilm assays were done as described previously (1,5). Cultures were grown overnight in TY broth and diluted 1:100 in BHIS-1% glucose-50 mM sodium phosphate buffer in a 24-well polystyrene plate. Plates were incubated statically for 24 hours. Supernatants were removed, then the adherent biofilms were washed once with PBS and stained for 30 minutes with 0.1% crystal violet. After the crystal violet was removed, the biofilms were washed once more with PBS and the remaining crystal violet was solubilized with 1 mL ethanol. Absorbance at 570 nm was measured.
