## Supplemental Data for "Multiple regulatory mechanisms control the production of CmrRST, an atypical signal transduction system in *Clostridioides difficile*"

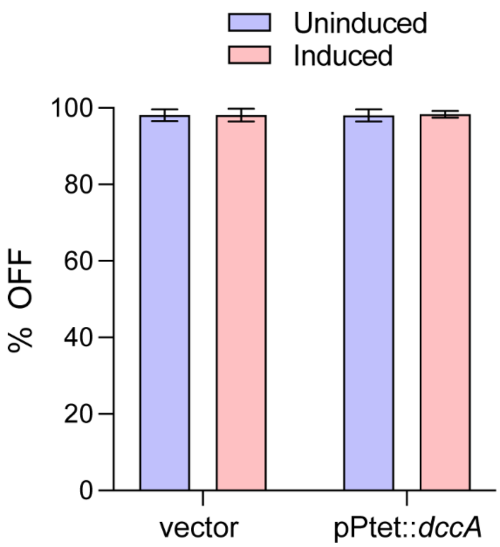


**Fig S1. C-di-GMP does not cause inversion of the *cmr* switch.** Δ*cmrR*::SNAP carrying $pP_{tet}$::*dccA* or vector control was grown to mid-exponential phase in BHIS broth with or without 20 ng/ml ATc. gDNA was collected for qPCR analysis of *cmr* switch orientation. Data are expressed as the percent OFF orientation. Shown are the means and standard deviations of six biological replicates from two independent experiments. No significant differences, two-way ANOVA with Tukey's multiple comparisons.

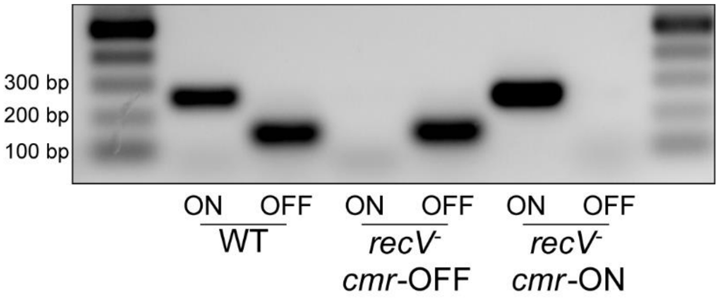


**Fig S2. The *cmr* switch is phase locked in *recV*-deficient strains.** Orientation-specific PCR to detect the orientation of the *cmr* switch in WT, *recV cmr*-OFF, and *recV cmr*-ON.

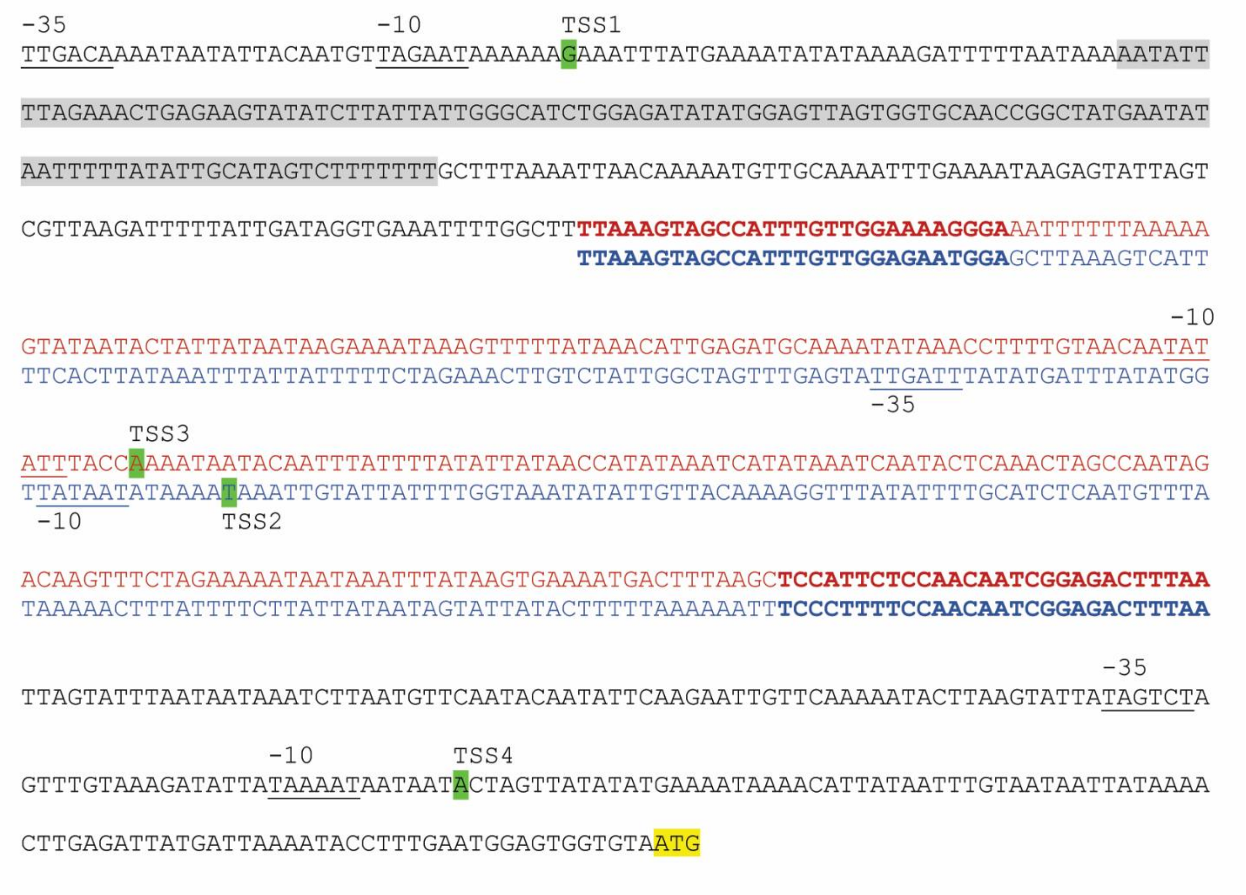


**Fig S3. Identification and mapping of multiple transcriptional start sites upstream of *cmrRST***. Map of transcriptional start sites (TSS) identified by 5' RACE in *recV cmr*-ON, *recV cmr*-OFF and *recV cmr*-OFF pDccA. Gray highlight indicates the Cdi-2-2 riboswitch sequence. Red and blue text denote the OFF and ON orientations of the *cmr* switch, respectively. Bold text indicates the inverted repeats. Green highlights mark TSS. Underlined text indicates putative -10/-35 sites. Yellow highlight denotes the *cmrR* start codon.

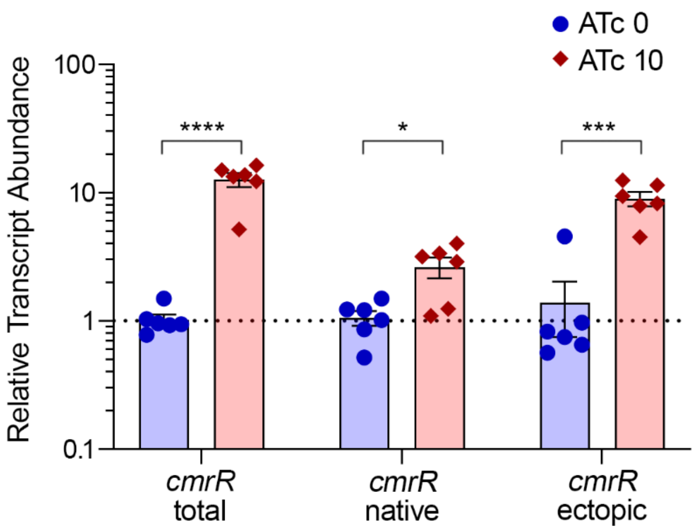


**Fig S4. Inducible expression of *cmrR* from an ectopic chromosomal site.** R20291 *recV* CDR2492::P$_{tet}$::*cmrR* was grown to mid-exponential phase in BHIS medium with and without 10 µg/mL ATc to induce *cmrR* expression. qRT-PCR was used to measure *cmrR* transcript abundance using 3 different primer sets to distinguish *cmrR* expression from the native site (primers R3111/R3112), from the ectopic site (primers R3113/R3112), and total *cmrR* (primers R2298/R2299). Data are expressed as relative transcript abundance in the ATc condition compared to that without ATc, with normalization to *rpoC* as the reference strain. Induction with ATc resulted in a 9.0-fold increase in *cmrR* mRNA from the ectopic site and a 2.6-fold increase from the native *cmrRST* locus, with a cumulative 12.7-fold increase in *cmrR* transcript. Shown are means and standard error. * $p < 0.05$, *** $p < 0.001$, **** $p < 0.0001$ by unpaired t-test.

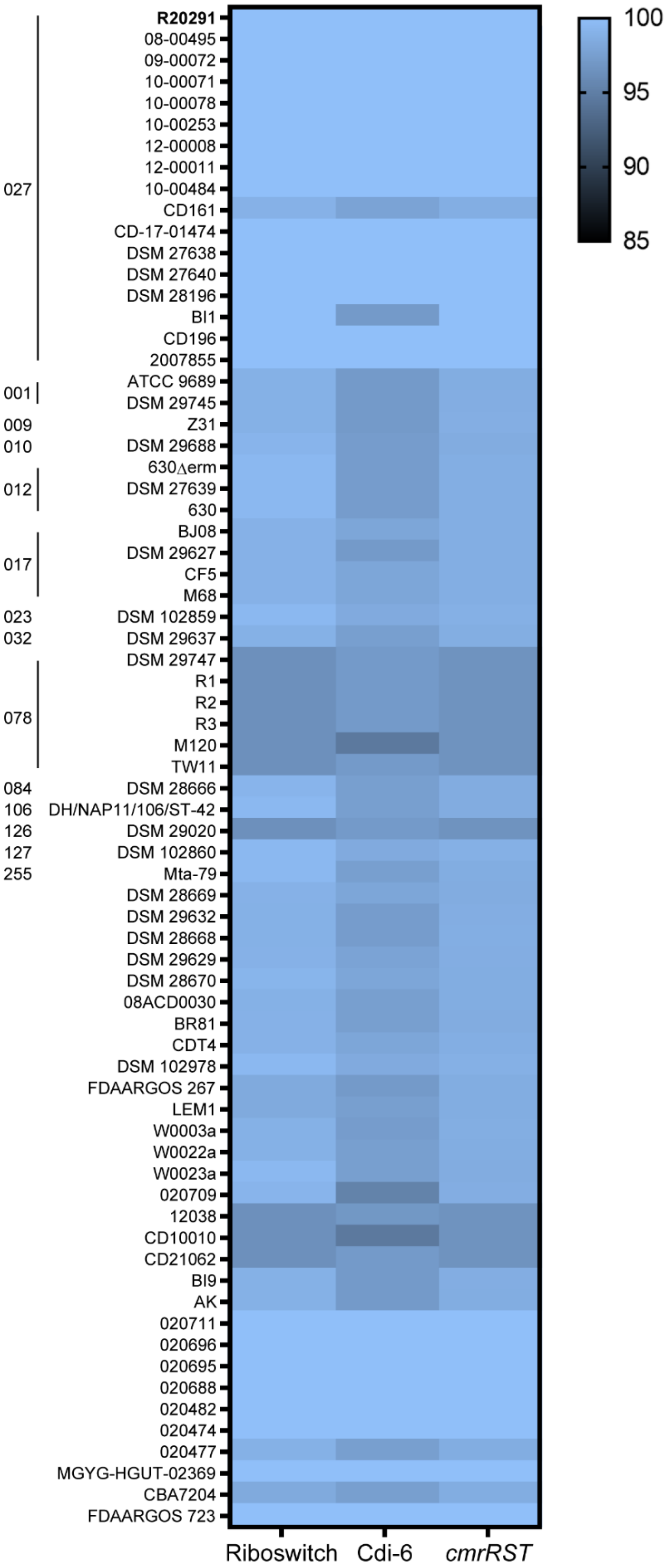


**Fig S5. *cmrRST* and upstream regulatory sequences are highly conserved across *C. difficile* strains and ribotypes.** Heat map showing percent sequence identity of 71 sequenced *C. difficile* strains in the NCBI database compared to the R20291 reference sequence (NCBI accession number FN545816.1). If the strain has been ribotyped, the ribotype is indicated on the left. For the *cmr* switch, both orientations were used as the reference and the highest percent sequence identity is shown. The following nucleotides were used as the reference sequence from R20291: riboswitch, 3,736,512-3,736,863; *cmr* switch, 3,735,969-3,736,513; *cmrRST*, 3,732,474-3,735,968.
