## Supplemental Tables for "Multiple regulatory mechanisms control the production of CmrRST, an atypical signal transduction system in *Clostridioides difficile*"

### S1 Table. Strains and plasmids used in this study

| Lab Notation | Strain Name | Description | Reference |
|---|---|---|---|
| AC472 | *Escherichia coli* DH5a | F- φ80*lacZ*ΔM15 Δ(*lacZYA*-*argF*)U169 *recA1 endA1 hsdR*17(rκ -, mκ+) *phoA supE44 thi*-1 *gyrA96 relA1* λ- *tonA* | Invitrogen (1) |
| RT270 | *Escherichia coli* HB101(pRK24) | *E. coli* used in conjugations with *C. difficile*, $Ap^R$, $Cm^R$ | (2) |
| RT273 | *C. difficile* R20291 | Ribotype 027 strain, WT (Genbank Accession # FN545816) | (3) |
| RT2395 | R20291 *cmr*-Δ3 OFF | R20291 with the *cmr* invertible element locked in the OFF orientation due to the deletion of three nucleotides in the right inverted repeat | This work |
| RT2406 | R20291 *cmr*-Δ3 ON | R20291 with the *cmr* invertible element locked in the ON orientation due to the deletion of three nucleotides in the right inverted repeat | This work |
| RT2256 | R20291 Δ*cmrR* | R20291 with in-frame deletion of *cmrR* | (4) |
| RT2296 | R20291 Δ*cmrR*Δ*cmrT* | R20291 with in-frame deletions of *cmrR* and *cmrT* | This work |
| RT2435 | R20291 *cmr*-Δ3 ON vector | R20291 *cmr*-ON with pRT1611; vector control | This work |
| RT2436 | R20291 *cmr*-Δ3 ON $pP_{tet::}$*dccA* | R20291 *cmr*-ON with pRT1587 for inducible dccA expression to increase c-di-GMP | This work |
| RT2437 | R20291 *cmr*-Δ3 ON $pP_{tet}$::EAL | R20291 *cmr*-ON with pRT2444 for inducible *pdcA* EAL domain expression to decrease c-di-GMP | This work |
| RT2438 | R20291 *cmr*-Δ3 OFF vector | R20291 *cmr*-OFF with pRT1611; vector control | This work |
| RT2439 | R20291 *cmr*-Δ3 OFF $pP_{tet}$::*dccA* | R20291 *cmr*-OFF with pRT1587 for inducible dccA expression to increase c-di-GMP | This work |
| RT2440 | R20291 *cmr*-Δ3 OFF $pP_{tet}$::EAL | R20291 *cmr* locked OFF with pRT2444 for inducible *pdcA* EAL domain expression to decrease c-di-GMP | This work |
| RT2187 | R20291 *cmrR*::SNAP | R20291 with *cmrR* replaced by allelic exchange with a SNAP-tag coding sequence | (5) |
| RT2500 | R20291 *cmrR*::SNAP vector | R20291 *cmrR*::SNAP with pRT1611 | This work |
| RT2501 | R20291 *cmrR*::SNAP $pP_{tet}$::*dccA* | R20291 *cmrR*::SNAP with pRT1587 for inducible dccA expression to increase c-di-GMP | This work |
| RT1693 | R20291 *recV cmr-OFF* | R20291 with an insertional mutation in *recV* (*recV::ermB)*; *cmr* locked OFF | (6) |
| RT2502 | R20291 *recV cmr*-OFF pMC123::*phoZ* | R20291 *recV::ermB cmr*-OFF with pMC123::*phoZ* (vector control) | This work |
| RT2507 | R20291 *recV cmr*-OFF pMC123::TSS4-*phoZ* | R20291 *recV::ermB cmr*-OFF with pRT2497 with reporter for TSS4 region only | This work |
| RT2516 | R20291 *recV cmr*-OFF pMC123::*cmr*OFF-*phoZ* | R20291 *recV::ermB cmr* OFF with pRT2514 with reporter for *cmr* OFF sequence | This work |
| RT2517 | R20291 *recV cmr*-OFF pMC123:: *cmr*ON-*phoZ* | R20291 *recV::ermB cmr* OFF with pRT2515 with reporter for *cmr* ON sequence | This work |
| RT1615 | R20291 vector | R20291 with pRT1611 (vector control) | (7) |

| | | | |
|---|---|---|---|
| RT2085 | R20291 pCmrR | R20291 with pRT2073 ($P_{tet}$::*cmrR*) | (4) |
| RT2107 | R20291 pCmrT | R20291 with pRT2106 ($P_{tet}$::*cmrT*) | (4) |
| RT2463 | R20291 *cmr*-Δ3 OFF pCmrR | R20291 *cmr* locked OFF with pRT2073 ($P_{tet}$::*cmrR*) | This work |
| RT2465 | R20291 *cmr*-Δ3 ON pCmrR | R20291 *cmr* locked ON with pRT2106 ($P_{tet}$::*cmrT*) | This work |
| RT2269 | R20291 Δ*cmrT* vector | R20291 Δ*cmrT* with pRT1611 (vector control) | (4) |
| RT2402 | R20291 Δ*cmrT* pCmrR | R20291 Δ*cmrT* with pRT2073 ($P_{tet}$::*cmrR*) | This work |
| RT2270 | R20291 Δ*cmrT* pCmrT | R20291 Δ*cmrT* with pRT2106 ($P_{tet}$::*cmrT*) | (4) |
| RT2183 | R20291 *recV cmr*-OFF pMC-Pcpr | R20291 *recV::ermB cmr* OFF with pMC-Pcpr | This work |
| RT2184 | R20291 *recV cmr*-OFF pDccA | R20291 *recV::ermB cmr* OFF with pMC-Pcpr::*dccA* | This work |
| RT1697 | R20291 *recV cmr*-OFF pRecV | R20291 *recV::ermB cmr* OFF with $pP_{tet}$-RecV | (8) |
| RT2520 | R20291 *recV cmr*-ON | R20291 *recV::ermB cmr* locked ON, derived from RT1693 | This work |
| RT2198 | R20291 *recV cmr*-OFF vector | R20291 *recV::ermB cmr* OFF with pRT1611 (vector control) | This work |
| RT2543 | R20291 *recV cmr*-OFF pCmrR | R20291 *recV::ermB cmr* OFF with pRT2073 ($P_{tet}$::*cmrR*) | This work |
| RT2544 | R20291 *recV cmr*-ON vector | R20291 *recV::ermB cmr* ON with pRT1611 (vector control) | This work |
| RT2545 | R20291 *recV cmr*-ON pCmrR | R20291 *recV::ermB cmr* ON with pRT2073 ($P_{tet}$::*cmrR*) | This work |
| RT2826 | R20291 *recV* CDR2492::$P_{tet}$::*cmrR* | R20291 *recV::erm* with ATc-inducible *cmrR* integrated between CDR20291_2492 and 2293 | This work |
| RT2827 | R20291 *recV* CDR2492::$P_{tet}$::*cmrR* vector | Inducible *cmrR* strain with pRT1343 (pMC123::*phoZ*) | This work |
| RT2828 | R20291 *recV* CDR2492::$P_{tet}$::*cmrR* pMC123::TSS4-*phoZ* | Inducible *cmrR* strain with pRT2497 | This work |
| RT2829 | R20291 *recV* CDR2492::$P_{tet}$::*cmrR* pMC123::*cmr*OFF/TSS4-*phoZ* | Inducible *cmrR* strain with pRT2514 | This work |
| RT2830 | R20291 *recV* CDR2492::$P_{tet}$::*cmrR* pMC123::*cmr*ON/TSS4-*phoZ* | Inducible *cmrR* strain with pRT2515 | This work |
| RT2831 | R20291 *recV* CDR2492::$P_{tet}$::*cmrR* pMC123::5'UTR*cmr*OFF-*phoZ* | Inducible *cmrR* strain with pRT2565 | This work |
| RT2832 | R20291 *recV* CDR2492::$P_{tet}$::*cmrR* pMC123::TSS1-*phoZ* | Inducible *cmrR* strain with pRT2566 | This work |
| RT2833 | R20291 *recV* CDR2492::$P_{tet}$::*cmrR* pMC123::*cmr*OFF-*phoZ* | Inducible *cmrR* strain with pRT2567 | This work |
| RT2834 | R20291 *recV* CDR2492::$P_{tet}$::*cmrR* pMC123::*cmr*ON-*phoZ* | Inducible *cmrR* strain with pRT2568 | This work |
| **Lab Notation** | **Plasmid Name** | **Description** | **Reference** |
| | pMTL-SC7215 | Vector for allelic exchange in *C. difficile* R20291 | (9) |
| | pRPF185 | *E. coli – C. difficile* shuttle vector, contains ATc-inducible $P_{tet}$ promoter with *gusA* | (10) |

| | | | |
|---|---|---|---|
| | pRT1611 | Derivative of pRPF185 with *gusA* removed, vector control | (7) |
| pRT1587 | pP$_{tet}$::*dccA* | pRPF185 with *gusA* replaced by *dccA*, ATc-inducible expression | This work |
| pRT2444 | pP$_{tet}$::EAL | pRPF185 with *gusA* replaced by EAL domain sequence from *pdcA*, ATc-inducible expression | This work |
| pRT2073 | pCmrR | pRPF185 with *gusA* replaced by *cmrR*, ATc-inducible expression | (4) |
| pRT2106 | pCmrT | pRPF185 with *gusA* replaced by *cmrT*, ATc-inducible expression | (4) |
| | pMC123 | *E. coli – C. difficile* shuttle vector | (2) |
| pRT402 | pDccA | pMC-Pcpr::*dccA* (nisin-inducible) | (11) |
| pRT1343 | pMC123*::phoZ* | pMC123 with *Enterococcus faecalis phoZ* | (7) |
| pRT2497 | pMC123*::*TSS4-*phoZ* | *phoZ* transcriptional reporter of the region between *cmrR* and right inverted repeat of the *cmr* invertible element | This work |
| pRT2514 | pMC123*::cmr*OFF/TSS4-*phoZ* | *phoZ* transcriptional reporter of the region from the *cmr* invertible element (OFF) to *cmrR* | This work |
| pRT2515 | pMC123*::cmr*ON/TSS4-*phoZ* | *phoZ* transcriptional reporter of the region from the *cmr* invertible element (ON) to *cmrR* | This work |
| pRT2566 | pMC123::TSS1-*phoZ* | phoZ transcriptional reporter of the region from the TSS1 promoter/c-di-GMP riboswitch to the LIR of the *cmr* switch | This work |
| pRT2567 | pMC123*::cmr*OFF-*phoZ* | *phoZ* transcriptional reporter of the region from the *cmr* invertible element (OFF) excluding TSS4 region | This work |
| pRT2568 | pMC123*::cmr*ON-*phoZ* | *phoZ* transcriptional reporter of the region from the *cmr* invertible element (ON) excluding TSS4 region | This work |
| pRT2565 | pMC123::5'UTR*cmr*OFF-*phoZ* | *phoZ* transcriptional reporter of the full *cmrRST* regulatory region with *cmr* switch OFF | This work |
| | pMSR0 | Vector for allelic exchange in *C. difficile* R20291, uses toxin-antitoxin counterselection | (12) |
| pRT2825 | pMSR0::CDR2493-P*tet*::*cmrR*-CDR2493 | Allelic exchange vector for inserting P*tet*::*cmrR* between CDR20291_2492 and CDR20291_2493 | This work |

**S2 Table. Primers used in this study.**

| Primer Name | Primer Sequence (5' to 3') |
|---|---|
| OS383 | GTTTTTTGTTACCCTAAGTTTAGTAATAGTATCAAGAGAAGAAGG |
| OS384 | AGTCTCCGGTTGGAGAATGGAGCTTAAAG |
| OS385 | TCTCCAACCGGAGACTTTAATTAGTATTTAATAATAAATC |
| OS386 | GATTATCAAAAAGGAGTTTCCATTACTTGCTCTAATAGAC |
| OS268 | TTTTTTGTTACCCTAAGTTTGCCATCCATGTATCCTATC |
| OS269 | TTAATATATTTATATAATCACTCCCAAATATCATTATTC |
| OS271 | AGATTATCAAAAAGGAGTTTTGTCAAATCTTGTAACCAAC |
| OS283 | AGTGATTATAGAAAATTATAACAATAAGAGGAGC |
| OS387 | GACTTTAAGCTCCATTCTCCAACAAT |
| OS137 | GAACAATTCTTGAATATTGTATTGAACATTAAGA |
| OS109 | GGAGATATATGGAGTTAGTGGTGCAA |
| OS111 | CGCTCTACTATATCCATAGCATCTTT |
| OS388 | AGTCTCCGGTTGGAAAAGGGAAATTTTTTAAAAAG |
| OS389 | TTTTCCAACCGGAGACTTTAATTAGTATTTAATAATAAATC |
| OS390 | CTTTTTAAAAAATTTCCCTTTTCCAACAAT |
| OS391 | CACCACTCCATTCAAAGGTATTTTAATC |
| R1907 | CAGAGCTCCTAGTACAAAGTATTTTATTTTGGAG |
| R1908 | GACGGATCCCAGTACTTAATGTCAATATCTTGTATAG |
| R2689 | CAGAGCTCCTATGGAGGAGATAAGTATATGGATTTAAAGGCATCAATTAATGATG |
| R2690 | CAGGATCCATACTATTCCCAATTTAACATCC |
| R2754 | ATGCAGAATTCACTTTAATTAGTATTTAATAATAAATCTTAATG |
| R2342 | ATGCAGGATCCCATTACACCACTCCATTCAAAGG |
| R2337 | ATGCAGAATTCGGTGAAATTTTGGCTTTTAAAGTAGCC |
| R2270 | GGAGATATATGGAGTTAGTGGTGCAA |
| R2271 | CTAGCCAATAGACAAGTTTCTAGAAAAATA |
| R2272 | GAACAATTCTTGAATATTGTATTGAACATTAAGA |
| R850 | CTAGCTGCTCCTATGTCTCACATC |
| R851 | CCAGTCTCTCCTGGATCAACTA |
| R2918 | CTCCTGAAATACTTGATGAACCAGAAGCTTAAGACCCACTTTCACATTTAAGTTG |
| R2919 | GGTATATGATATGAAAGAGAGAGTCTCAAACAGTATCTCACTTATGGTACAAACTTATATCC |
| R2914 | CATTGATTTCTTTCAGTTTCGGATCCCTCCTGAAATACTTGATGAACCAGAAG |
| R2915 | CAACTTAAATGTGAAAGTGGGTCTTAAGGTGAACTTTTGTTCATGAGACACTC |
| R2916 | GGATATAAGTTTGTACCATAAGTGAGATACTGTTTGAGACTCTCTCTTTCATATCATATACC |
| R2917 | GACGTCGACTCTAGAGGATCCTACATGCAATAACAGTACCTCAGGA |
| R2743 | GTGTTATCAATTGCACTACTCATGG |
| R2744 | GTTGAACCATTAGCTAAGGATTCAG |
| R2987 | GTATGTTAGAAGTTGTTTCAGAAGGC |
| R2988 | GTGGCTGAAGTAGTATCAGAAGC |

| | |
|---|---|
| R2298 | AAGAAAAAGTTCGGGGATTTTTAGC |
| R2299 | CGCTGAAAACTTTAACACATTAGGA |
| R2537 | GACAAGGATAATTGCC |
| R2538 | CCATCACCATCAGTTAG |
| R2539 | GATAGATGACTGGGAG |
| R2540 | CGATAAGTAGCATTCCC |
| R2745 | CTTTTTTTGCTTTAAAATTAACAAAAATGTTGC |
| R2746 | CTTTAAAAGCCAAAATTTCACCTATCAATA |
| R2751 | CTTAATGTTCAATACAATATTCAAGAATTG |
| R2803 | GAAAATAAAACATTATAATTTGTAATAATTATAAAACTTGAG |
| R2804 | CCAGTAAATTATACATTACACCACTCC |
| R2716 | GTCTCTGATGTATATACATTAAAACC |
| R2273 | TCATTACCAGGTGTAGCAGTGAATGC |
| R2274 | GATAGAGCATGGTCCTTGAGCTTCT |
| R2792 | GTCTAAAACTATACGCTC |
| R2793 | CCATAGCATCTTTAGCAGTCTCTGATGTATATA |

Restriction sites are underlined. Regions of homology for Gibson assembly are lowercase.
